# BioStructNet: Structure-Based Network with Transfer Learning for Predicting Biocatalyst Functions

**DOI:** 10.1101/2024.10.16.618725

**Authors:** Xiangwen Wang, Jiahui Zhou, Jane Mueller, Derek Quinn, Thomas S. Moody, Meilan Huang

**Affiliations:** School of Chemistry and Chemical Engineering, Queen’s University Belfast, BT9 5AG, Northern Ireland, U.K.; Department of Biocatalysis and Isotope Chemistry, Almac Sciences, Craigavon, BT63 5QD, Northern Ireland, U.K.; Arran Chemical Company Limited, Unit 1 Monksland Industrial Estate, Athlone, Co. Roscommon, N37 DN24, Ireland

## Abstract

Enzyme-substrate interactions are essential to both biological processes and industrial applications. Advanced machine learning techniques have significantly accelerated biocatalysis research, revolutionizing the prediction of biocatalytic activities and facilitating the discovery of novel biocatalysts. However, the limited availability of data for specific enzyme functions, such as conversion efficiency and stereoselectivity, presents challenges for prediction accuracy. In this study, we developed BioStructNet, a structure-based deep learning network that integrates both protein and ligand structural data to capture the complexity of enzyme-substrate interactions. Benchmarking studies with the different algorithms showed the enhanced predictive accuracy of BioStructNet. To further optimize the prediction accuracy for the small dataset, we implemented transfer learning in the framework, training a source model on a large dataset and fine-tuning it on a small, function-specific dataset, using the CalB dataset as a case study. The model performance was validated by comparing the attention heat maps generated by the BioStructNet interaction module, with the enzyme substrate interactions revealed by enzyme-substrate complexes revealed from molecular simulations. BioStructNet would accelerate the discovery of functional enzymes for industrial use, particularly in cases where the training datasets for machine learning are small.

## Introduction

Compound-protein interactions (CPIs) play a crucial role in the design of enzyme as biocatalysts, and the accurate and fast prediction of CPIs can drive innovations in biocatalysis and related fields. Although experimental assays remain the most reliable approach for determining CPIs, the vast costs and labor required to experimentally characterize every potential protein-ligand pair make this approach prohibitive. In recent years, computational methods, particularly machine learning techniques, have greatly advanced the CPI prediction.^1,2^ Compared to the deep-learning based protein structure prediction, for which there are already established datasets, i.e. CASP,^3,4^ little has been reported for assessing the biocatalysis properties predicted by machine learning. The lack of standardized protocols and benchmark data sets, coupled with the discrepancies in the performance measurements, presents significant challenges in developing accurate predictive models in the enzyme catalysis arena.^5^ Moreover, most ML models showed limited predictive accuracy due to overfitting when applied to small, function-specific datasets, especially for those protein sequences with high similarity. Consequently, there is an urgent need for innovative methodologies that can effectively leverage limited function-specific enzyme data by applying the insight gained from large datasets for improving the prediction accuracy for small datasets.

As powerful tools for CPI prediction, machine learning algorithms, utilize large datasets to uncover complex patterns,^6^ using various techniques such as random forest (RF),^7^ support vector machine (SVM),^8^ Gaussian Processes,^9^ and boosting.^10^ In recent years, deep learning algorithms such as convolutional neural networks (CNNs) and graph neural networks (GNNs) have proven effective in predicting enzyme functions based on the features retrieved from protein sequences.^11–20^ With the recent increase in protein structural data and protein-ligand complexes datasets, structure-based machine learning methods^21–24^ have been reported, where proteins are represented through sequential CNNs or 2D pairwise distance maps.

Despite the emerging machine learning models for CPI prediction, these models have limited performance in predictive accuracy when applied to small, function-specific datasets due to overfitting and the lack of diverse training examples.^20^ This limitation necessities the development of novel methods that can effectively leverage the available data while mitigating the impact of dataset size. Transfer learning, which involves pre-training a model on a large dataset and fine-tuning it on a smaller, task-specific dataset, offers a promising solution.^25,26^ By transferring knowledge from a broad, general model to a specialized model, transfer learning can enhance the prediction accuracy for function-specific enzyme-substrate interactions.

Compared to machine learning methods, physics-based computational approaches such as molecular docking and molecular dynamics (MD) simulations are also used for CPI prediction. However, careful analysis is required when using these methods due to the challenges in conformational sampling. Deep learning was integrated with molecular docking results in drug discovery, where MD simulated trajectories were employed as the input features of machine learning models to improve the prediction accuracy^27,28^ or enhancing the efficiency of traditional docking-based screening. ^29,30^ Enlighted by these advances, we propose to concinnate the data-based parameters with physics-based computational simulations to validate and interpret the deep learning model’s predictions. This approach could be used to assess the reliability and accuracy of the deep learning predictors in the context of enzyme-substrate interactions.

One enzyme of particular interest for CPI prediction and application is Candida antarctica lipase B (CalB).^31^ In recent years, the rapid increase in plastic waste has highlighted the need for innovative solutions to degrade plastics efficiently. CalB has emerged as a promising biocatalyst due to its high thermostability and effectiveness in breaking down polyethylene terephthalate (PET) into terephthalic acid (TPA), a key step in plastic degradation. ^32^ Moreover, CalB can degrade other plastics, such as *ε* -polycaprolactone, further underscoring its potential in environmental applications. ^33^ CalB belongs to EC3 hydrolase class, which hydrolyzes the ester bond in a large and promiscuous binding pocket with an oxyanion hole, represent a challenging case for structure-based machine learning. The growing importance of CalB in plastic degradation and its challenging structural and mechanism features, makes it an ideal case study for developing and validating a robust CPI prediction model, especially for small and function-specific datasets.

In this study, we developed a novel structure-based deep learning model BioStructNet to address the challenges of small, function-specific enzyme datasets, using CalB as a case study. The CalB database contains a limited number of sequences derived from the wildtype protein, therefore presenting a challenging case for developing machine learning models for CPI prediction. To address this challenge, we developed BioStructNet, utilizing transfer learning to transfer the knowledge learned from larger datasets, to enhance the model’s ability to generalize in small datasets like CalB. Protein graphs are generated based on residues coordinates and assigns physicochemical properties property features assigned to nodes, while ligand graphs are created from SMILES. These structure-based graph representations are combined in the interaction module to generate contact heat maps for specific protein-ligand pairs, which are then used for regression or classification predictions. By combining structural representations with transfer learning, BioStructNet significantly improves the prediction accuracy for CalB and similar small datasets. The high-scoring residues predicted by the deep learning model’s attention weights are in agreement with the key protein residue sites identified through docking and MD simulations, which validated the model’s reliability. Our BioStructNet transfer learning approach provides a promising solution to improve CPI prediction accuracy for small, specific datasets, and would accelerate the discovery of effective biocatalysts for specific functions.

## Results and Discussion

### BioStructNet framework

CalB belongs to the hydrolase enzyme family, which hydrolyzes ester substrates through reactions with water. Despite its significant industrial importance, the limited availability of data makes it difficult to build accurate deep learning models for CalB. To address this challenge, we designed a BioStructNet framework (Figure 1) to predict the function of small target datasets by leveraging the knowledge learned from the large source datasets.

**Figure 1:**
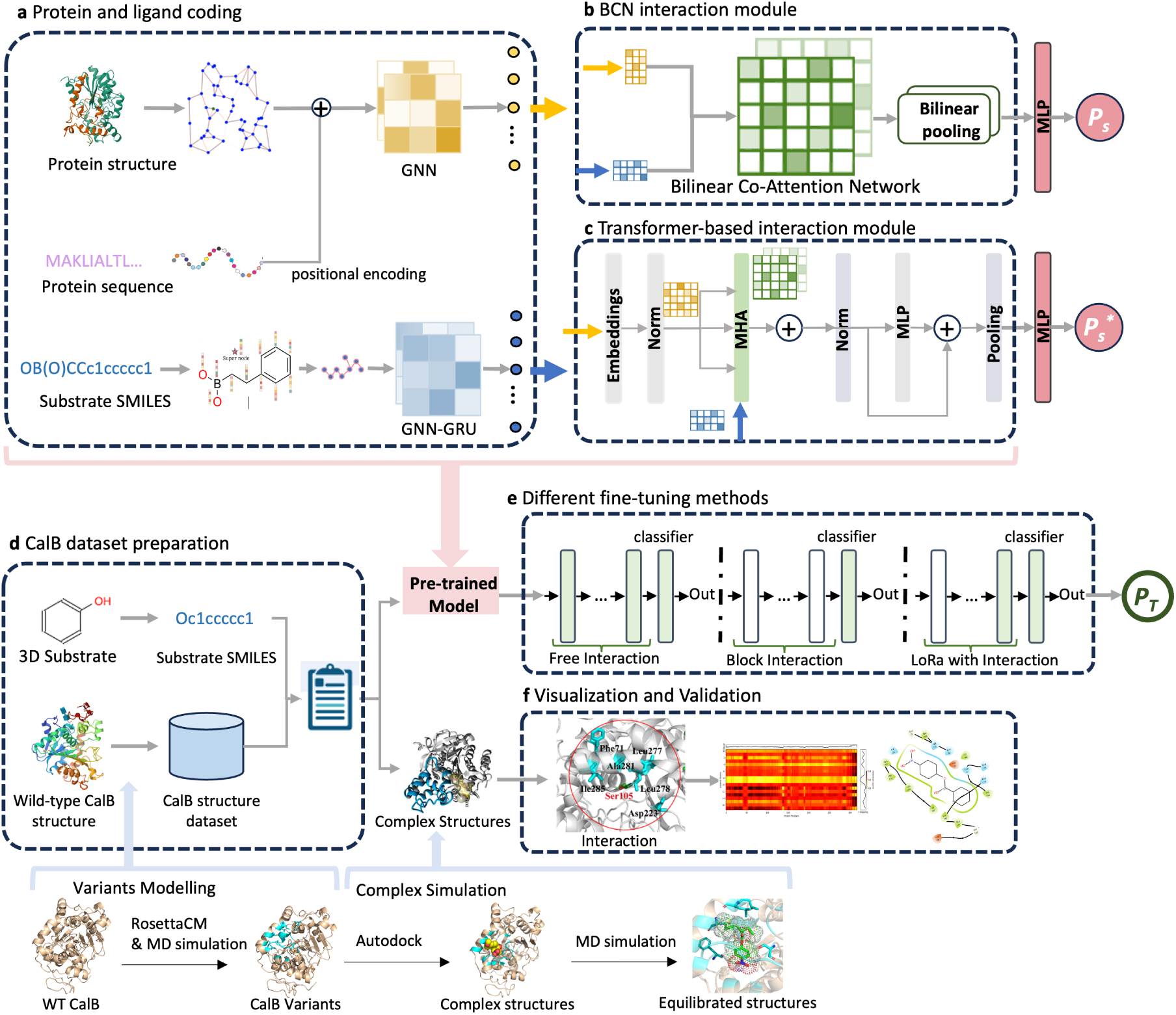
BioStructNet framework with two sections: processing the source task dataset (a, b, c) and transfer-learning and validation (d, e, f). **a.** Protein and Ligand Embeddings. Protein and ligands are encoded by GNNs, with substrate structures encoded by SMILES and protein structures denoted by 3D contact maps and sequence-based positional embeddings, to preserve spatial relationships. **b.** BCN Interaction Module. The Bilinear Co-Attention Network utilizes two fully connected networks to linearly transform the input features of proteins and ligands, followed by a bilinear multiplication to compute the attention weights. **c.** Transformer-based Interaction Module. This module with multi-head self-attention on embedded protein features preceding a cross-attention with ligand features, is embedded with layer normalization and implemented in a feed-forward network. **d.** CalB Data Preparation. Protein structures of CalB variants generated by RosettaCM and refined by MD simulations. Autodock is used to dock the CalB variants with ligands to give complex structures, which are then analyzed by MD simulations to evaluate interactions. **e.** Different fine-tuning strategies, including free interaction, block interaction, and LoRa for model adaptation. **f.** Visualization of protein-ligand interactions and attention weights predicted by BioStructNet.

The BioStructNet framework is divided into two main sections: modules for processing the source task dataset (blocks a, b, and c); and modules for transfer learning and validation (blocks d, e, and f). Protein structures are encoded using contact-map graphs, which are comprised of edges represented by the spatial distance between the alpha carbons of two residues within a defined distance cutoff and node features assigned with residue properties.^34^ Simultaneously, ligand structures are encoded by graphs comprised of edges and nodes represented by atomic bonds and atom types (Figure 1a). Attention mechanisms are employed using a bilinear co-attention network to capture the local-range CPIs information and a transformer-based interaction module, which integrates the global protein feature by multihead attention (MHA) among protein amino acids, to imply the long-range CPIs (Figure 1b and 1c). The transfer learning module employs three fine-tuning methods: a free interaction module, where all layers interact freely, allowing flexibility in adapting to new data; a block interaction module, where interaction layers are selectively constrained to preserve the structure of the pre-trained model; and a LoRa fine-tuning method, which combines low-rank adaptation (LoRa) with structured interactions to balance model efficiency and flexibility (Figure 1e). The training process utilizes 5-fold cross-validation and bootstrapping to ensure prediction accuracy.

The source model is built on a source database of hydrolases with the turnover number (kcat). The model performance for regression tasks is evaluated on its predictive accuracy for enzymes activity for substrates reflected by enzymes’ catalytic efficiency. Additionally, the predictive capability of the model for classification tasks is demonstrated using a widely applied human compound-protein interaction (CPI) dataset, which allows to differentiation of enzymes with distinct activities level reflecting different enzyme-substrate binding affinities. The target model is built on a CalB conversion database. The protein structures of CalB variants generated by comparative modelling using RosettaCM and refined by molecular dynamics (MD) simulations, serve as the input for the transfer learning. For model validation, the complex structures obtained by docking followed by MD simulations, were inspected to assess the protein-ligand interactions in relation to the attention weights of the model.

### BioStructNet Performance on the Source Datasets

The performance of BioStructNet was evaluated on regression task with Kcat “EC 3” (hydrolase) dataset and classification tasks with Human CPI dataset, in comparison to various baseline models, including random forest (RF), k-nearest neighbors (KNN), L2, Tsubaki’s^35^, DLKcat^19^, DrugVQA^21^, TransformerCPI2.0^36^, ALDELE^20^ and DrugBAN^16^ (Table 1). For the regression task, RMSE and R² were used as the metrics of model performance, while for the classification task, AUC, recall, and precision were used.

**Table 1:**
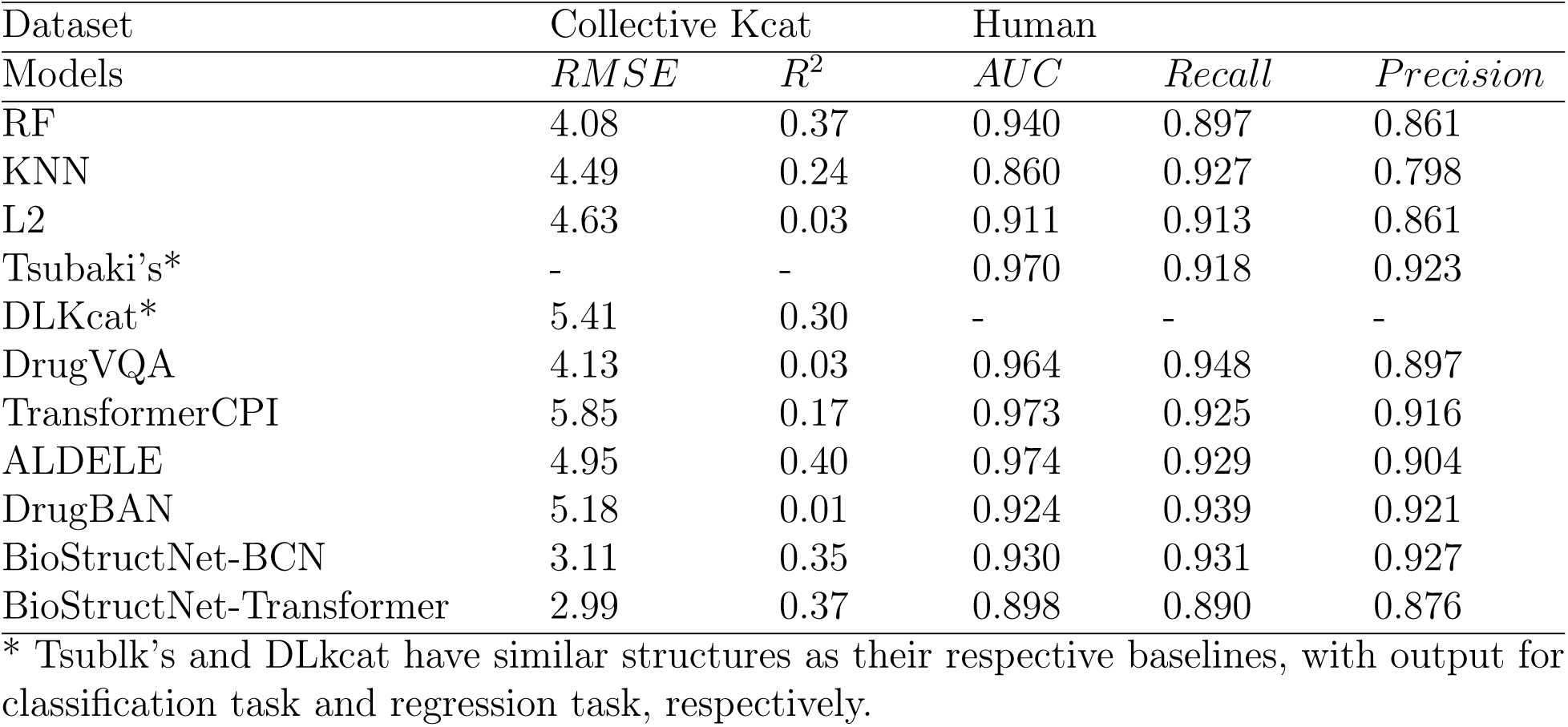
Benchmark studies of the performance of machine learning models in regression and classification tasks, using the Kcat “EC 3” dataset (for regression task) and the human dataset (for classification task), respectively.

In the regression task (Kcat dataset), 5-fold cross-validation was performed by splitting the data into training, validation, and test sets with a ratio of 7:1.5:1.5. The final model was selected based on the best R² score on the validation set, and performance metrics were averaged across the 5 folds to ensure robustness. Our BioStructNet model, with both BCN and Transformer interaction modules, achieved better RMSE (2.99 and 3.11) and R² scores compared to the baseline models, indicating improved performance in predicting Kcat values. We also benchmarked our BioStructNet with other approaches for handling classification tasks (predicting interaction or non-interaction, represented as 0 or 1), using the widely used human CPI dataset. Among all, the sequence-based Models performed particularly well (AUC around 0.97), such as ALDELE, TransformerCPI, and Tsubaki’s. These models are exclusively based on sequence information, hence they can be utilized to train larger datasets and demonstrate obvious advantages in terms of data coverage, as reflected in their high AUC, recall, and precision scores. Notably, our BioStructNet models, particularly the BCN interaction module, show competitive precision and recall scores. These results demonstrate the potential of BioStructNet’s for specific biocatalysis applications, particularly for those scenarios where the goal is to assure the predictive accuracy while minimizing the false positives.

The BioStructNet models trained with the Kcat ’EC 3’ dataset for the source task serves as the foundation for the CalB task with performance shown in Figure 2. The RMSE and R² values for the validation set during the training process are shown in Figure 2(a), and the performance of the final BioStructNet models—both with the BCN interaction module and the Transformer-based interaction module are shown in Figure 2(b). BioStructNet-BCN model displayed a RMSE of 3.11 and a R2 of 0.35. The BioStructNet-Transformer model showed a slight advantage over the BCN module with the RMSE of 2.99 and the R2 of 0.37. Because it implements a self-attention for protein information prior to the cross-attention with ligand, thus it is able to effectively capture global contexts and long-range dependencies in protein-ligand interactions. Therefore, to improve prediction accuracy and generalization, it is necessary to accurately represent complex interactions, by capturing both local and global interaction patterns using structural features encoded in graph-neural network.

**Figure 2:**
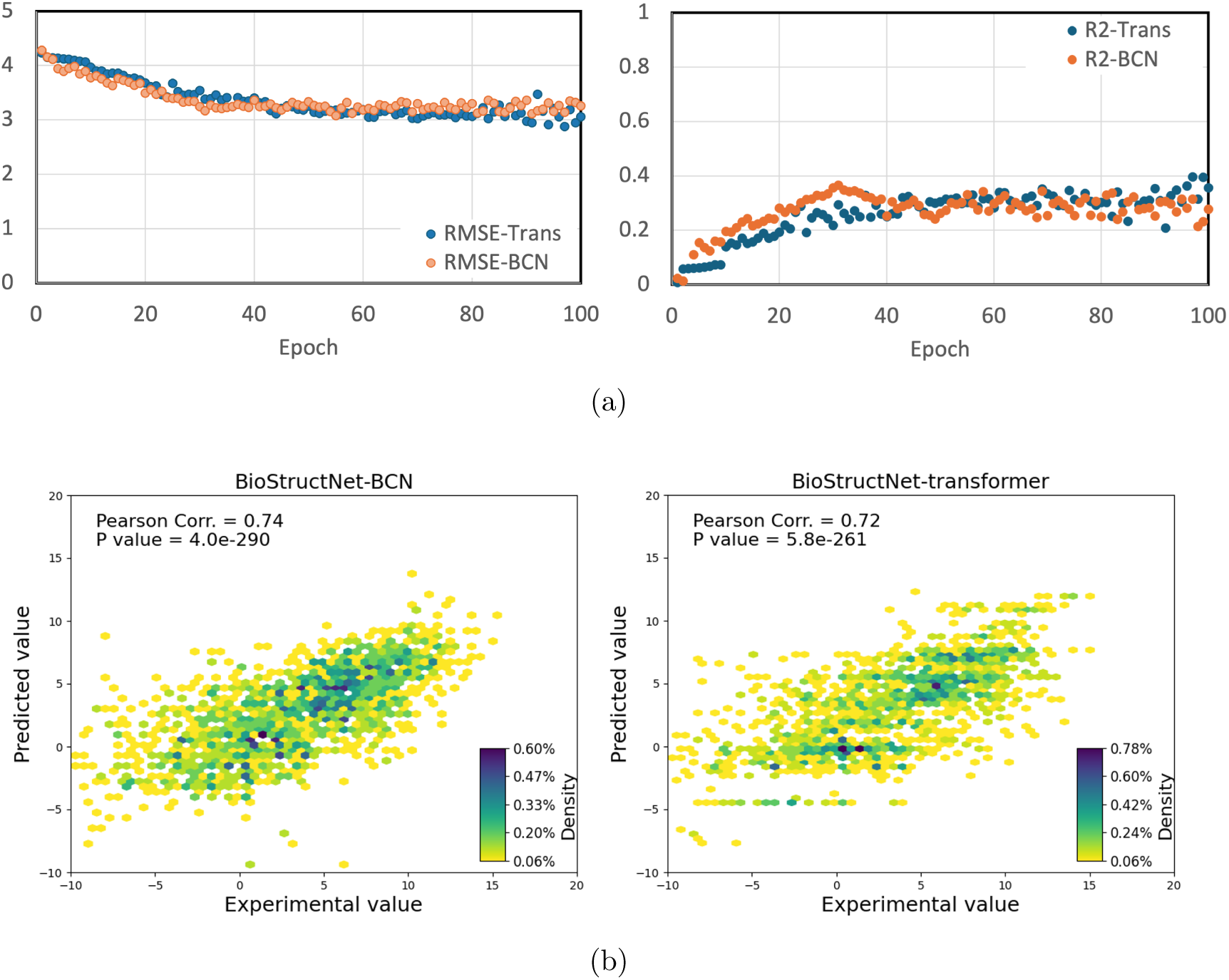
The performance of BioStructNet models for Kcat prediction with BCN interaction module and transformer-based interaction module. **a.** The r.m.s.e and r2 of validation set during the training process. **b.** Performance of the final deep learning models. The correlation between predicted Kcat values and the experimental values in the whole dataset (training, validation and test datasets) was evaluated. The brightness of colour represents the density of data points. P values for Pearson’s correlation were calculated.

### BiostructNet on the Target Dataset

The BiostructNet models are trained on a conversion dataset of CalB variants derived from the wild-type (WT) enzyme. The performance of the models was evaluated on the target CalB dataset for both regression and classification tasks, with a focus on implementing transfer learning in classification to improve prediction accuracy for small datasets.

First, the model was trained for the regression task and the performance is shown in Table 2. As expected, the results are inferior to those that are trained based on the large hydrolase Kcat dataset (Table 1). Among the tested models, BioStructNet-BCN exhibited the best performance with an RMSE of 9.31 and an R² of 0.37, while the BioStructNet- Transformer performed moderately with an RMSE of 12.18 and an R² of 0.09. Unlike the models trained on the large Kcat dataset, where the BioStructNet-Transformer interaction module performed better, the BCN interaction module shows better performance on the small CalB conversion dataset. This is because the variants in the CalB dataset are derived from the same wild type (WT) so that they share similar structures, and thus the attention mechanism of BCN is effective in capturing the subtle local interactions for the CalB dataset with high structural similarities. The transformer-based interaction module with a global self-attention mechanism incorporates long-range structural features, hence might not be as effective in the case of CalB where local differences are more critical. Thus, the BCN interaction module was specifically used to fine-tune the source model on the CalB dataset. In view of the deficient performance shown in the regression task for the CalB dataset, classification approach was explored to improve the prediction accuracy. A categorical binary classification approach with multiple thresholds was evaluated. Three fine-tuning methods—free, block, and LoRa—were systematically applied to the BCN interaction module. The transfer learning performance on the CalB conversion dataset are evaluated by AUC, Accuracy and RE (relative error), using three different classification thresholds: 15%, 30%, and 40% (Table 3). The 15% cutoff were applied to distinguish CalB variants with low conversion rates from moderate or high rates, to rule out the least active enzymes from the subsequent experimental validations. The 30% cutoff serves as an intermediate threshold, marking the variants with moderate enzyme activities, while the 40% cutoff distinguishes the variants with relatively high conversion rates, labelling the enzymes with exceptionally high conversion. The performance of BioStructNet transfer model are averaged across 100 bootstrapping iterations. The fine-tuning results for the transformer-based module for the source dataset are provided in the Supplementary Table S1.

**Table 2:**
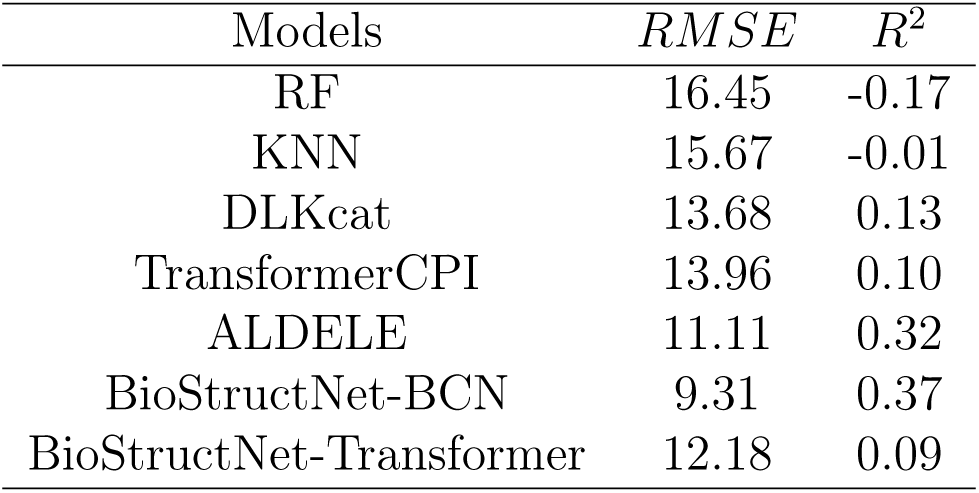
Performance comparison on the CalB conversion dataset for regression task.

| Models | $RMSE$ | $R^2$ |
| --- | --- | --- |
| RF | 16.45 | -0.17 |
| KNN | 15.67 | -0.01 |
| DLKcat | 13.68 | 0.13 |
| TransformerCPI | 13.96 | 0.10 |
| ALDELE | 11.11 | 0.32 |
| BioStructNet-BCN | 9.31 | 0.37 |
| BioStructNet-Transformer | 12.18 | 0.09 |

**Table 3:** The performance of transfer learning models fine-tuned on the BCN interaction module was evaluated on the CalB conversion dataset for the classification task.

| Cutoff | Models | AUC | Accuracy | 1.0% RE |
| --- | --- | --- | --- | --- |
| 15 | Free BCN | 0.63 | 81.38% | 0.28 |
|  | Block BCN | 0.71 | 82.47% | 0.30 |
|  | LoRa BCN | 0.71 | 83.41% | 0.31 |
| 30 | Free BCN | 0.64 | 66.27% | 0.45 |
|  | Block BCN | 0.68 | 68.85% | 0.43 |
|  | LoRa BCN | 0.69 | 72.71% | 0.47 |
| 40 | Free BCN | 0.66 | 62.62% | 0.48 |
|  | Block BCN | 0.69 | 65.76% | 0.44 |
|  | LoRa BCN | 0.66 | 68.17% | 0.48 |

The results indicate that the LoRa BCN model consistently outperforms other methods across all thresholds. At the 15% cutoff, LoRa BCN achieves the highest accuracy (83.41%) and a strong AUC (0.71), effectively distinguishing the CalB variants with low conversion rates from moderate and high rates. The accuracy of the fine-tuning model with 30% cutoff and the 40% cutoff, are inferior to that with the 15% threshold. This indicates the classification threshold would affect the model performance and for the CalB conversion dataset, a low cutoff value would be valuable to exclude the non-functional variants and hence reduce the variant library size for subsequent experimental evaluation.

The CalB dataset includes the conversion data of wild-type and variant proteins, for various substrates. In addition to the above classification with different thresholds, numerical binary classification is made based on the relative conversion value of a mutant’s compared with that of the wild-type protein for the same substrate: i.e. if the mutant performs comparably or better (taking into consideration of the experimental errors), it is classified as 1; otherwise, it is classified as 0. This method helps identify functional differences and informs beneficial mutation engineering. The CalB dataset is small with mutation positions populated only around several positions. Such small datasets can lead to performance variability. To improve the model’s robustness, we implemented a data augmentation strategy grounded in domain-specific knowledge of CalB enzyme catalysis. Considering mutations near active site residues around the catalytic triad Asp 187, Ser 105, and His 224 (1st and 2nd sphere regions)^37^ are more likely to affect the enzyme function than the mutations farther away, we generated 1,208 single-point mutation variants far away from the catalytic core (20 Å beyond the catalytic triad) and labelled them as the negative samples. This strategy expanded the CalB database to a much larger one with 145 positive and 1,298 negative samples (shown in Figure 3(a)). This strategy ensured both diversity and balance in the dataset, improving support for machine learning training and better generalization in the model.

**Figure 3:**
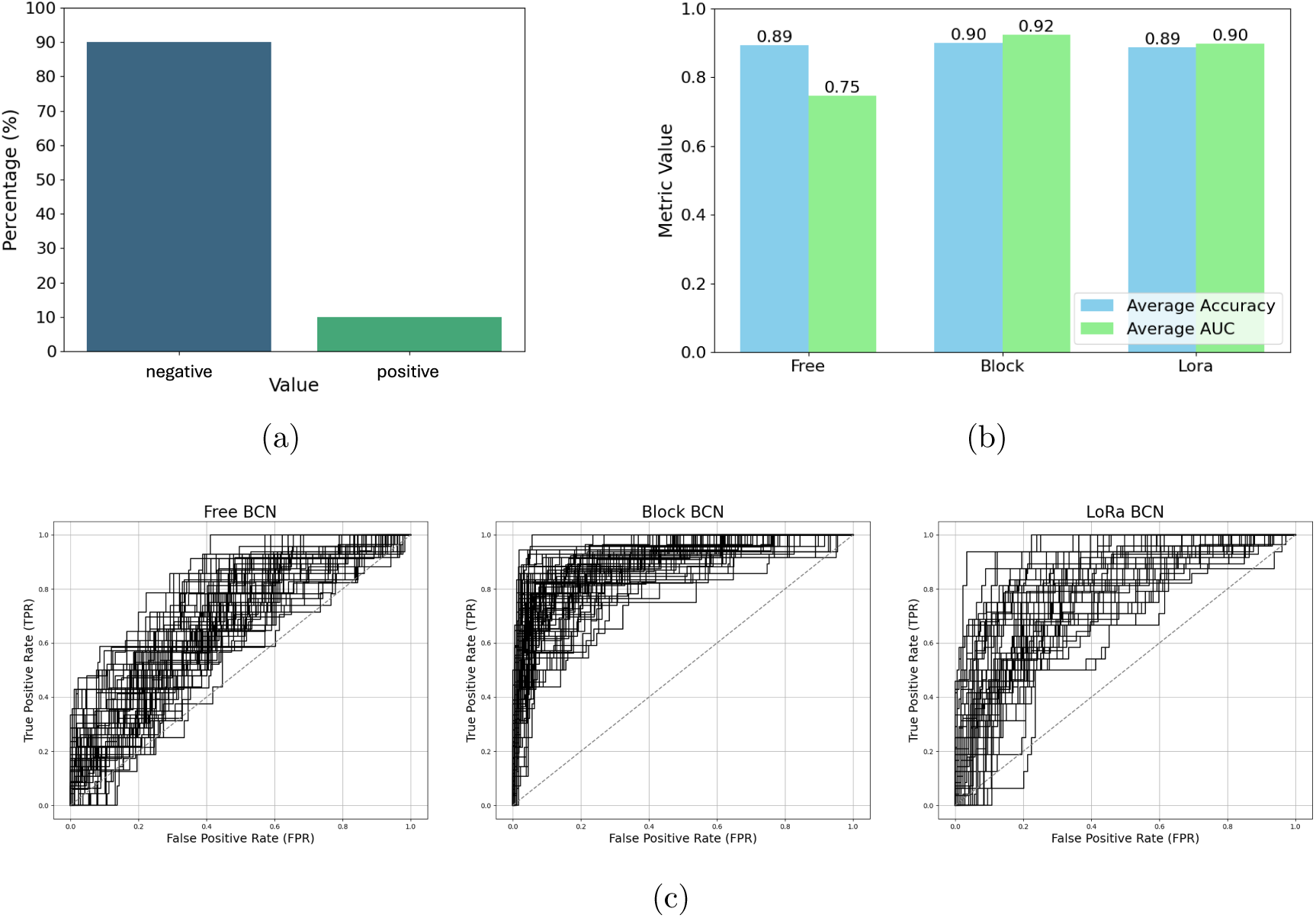
The performance of BioStructNet-BCN transfer learning models on the CalB enzyme dataset, which was classified based on the relative of the variants in comparison with the wild-type and an augmentation approach. a. Percentage distribution (0 and 1) of the augmented dataset. Approximately 90% of the values are 0, while around 10% of the values are 1. b. Comparison of Average Accuracy and AUC for the three fine-tuning strategies (Free, Block, and LoRa). c. The ROC curves for the final-epoch results of each boosting iteration, showing the FPR and TPR across the three fine-tuning methods: Free, Block and LoRA. Each solid black line represents the ROC for a boosting iteration, while the dashed black line shows the random classification.

The comparison of model performance metrics, accuracy and AUC (Figure 3(b)) and the receiving operating characteristic (ROC) curves across three fine-tuning methods—Free, Block, and LoRa shows the trade-offs between parameter flexibility and generalization in transfer learning (Figure 3(c)). The average AUC and accuracy for all methods surpass models with different conversion thresholds. with an accuracy of 90% and AUC of 0.90, indicating the superior model performance of the binary classification compared to the classification with definite thresholds (Table 3).

The BioStructNet with transfer learning achieved good performance for predicting the conversion of CalB regardless of the classification approaches, either categorical or numerical binary, effectively transferring the enzyme-substrate interaction information learned from the Kcat source dataset to the CalB conversion prediction. Due to the high sequence similarity in the target dataset, the diversity among different samples in the dataset are minimal, limiting the model’s ability to learn sufficient diversity. Therefore, leveraging the broad enzyme-substrate interaction information from the source Kcat dataset, which encompasses the diversity and complexity of protein binding with various small molecules.

Free fine-tuning, which adjusts all interaction parameters freely, offers greater flexibility during parameter optimization, however, it would result in noise and overfitting particularly for the small target dataset, as shown by the scattered ROC results and fluctuations at low FPR (Figure 3). In contrast, Block fine-tuning, which constrains the majority parameters in the original network, only adjusting the classifier and the final layer of the network during optimisation, retaining interaction information from the Kcat source dataset, and hence achieving high accuracy with a steep ROC curve. LoRa fine-tuning employs low-rank adjustments to fine-tune local network, retaining the performance of pre-trained model on the source dataset while adapting for the new target dataset by minimizing the parameter adjustment, as demonstrated by a stable ROC curve. Thus, it is crucial to select appropriate fine-tuning strategy, and the choice of fine-tuning methods should be guided by the prediction tasks for specific target dataset.

Overall, the BioStructNet framework with transfer learning approaches, with its ability to capture local protein-ligand interaction patterns by BCN interaction module, shows robust performance across different classification thresholds in the classification tasks for small function-based dataset such as the CalB conversion dataset.

### Model validation and attention visualization

For enzyme engineering, it is critical to identify the amino acid residues and interaction pairs of compound-protein complexes responsible for the catalytic activities of enzymes. BioStructNet’s attention mechanism captures the importance of features of interactions between protein residues and ligand atoms in the CPIs. The atom pairs with higher attention weights denote stronger or more dominant CPIs ascribed to both special distances and the physiochemical properties of amino acids. The protein-ligand interactions concentrate on the spatially close interacting pairs, so the atom pairs in the first sphere of the catalytic site tend to have higher attention weights.

To validate the predicted CPIs responsible for the conversion disclosed by the machine learning models, we investigated the binding modes of WT and mutant CalB enzymes using docking and MD simulations. Principal Component Analysis (PCA) of the equilibrated MD trajectories as demonstrated by the RMSD analysis, was performed to reduce the data into principal components (Supplementary Figure S3). K-means clustering was applied to identify the dominant conformations of the protein-ligand complexes, which were then analysed in relation to the attention weights from BioStructNet models (Supplementary Figure S4).

The distribution of attention weights for the CPIs between the wild-type enzyme with 25 various ligands is summarised in Figure 4 at the eight specific residues experimentally selected for mutation engineering. ^38^ The attention weights are highly concentrated between 0% and 20% for residues T42, S105 and A282, indicating that these positions are critical for interactions with the 25 different ligands involved in the study. The attention weights for residues S47 and W104 are primarily concentrated around 20%, although slightly spanning to top 40%, indicating diverse interaction potentials across different CPIs. Two hydrophobic residues I189 and V190 exhibit a wide range of attention weights, reflecting significant variability in the effect of mutations across different interactions. The attention weights for residue A281 are predominantly concentrated between 20% and 60%, with a noticeable peak around 50%.

**Figure 4:**
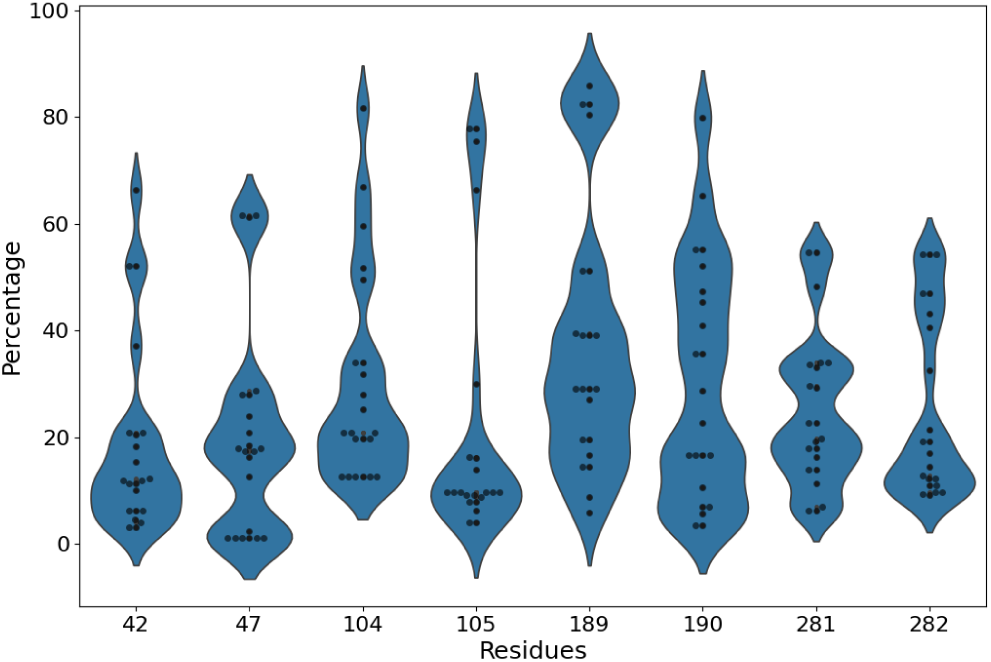
Violin plots of the attention weight distribution, representing different protein residues of wild-type CalB interacting with 25 various ligands. Wider regions indicate higher attention density at specific residue positions, and the wide lower bout of the violin indicates that attention weight is largely concentrated in the top percentage.

Here, the CPI between the wild-type protein and ligand rac-10 was shown as an example.^38^ The attention scores generated by BioStructNet were normalized to a 0-1 range to ensure consistency in the representations. The attention maps were averaged across all heads to obtain a single attention score matrix between ligand atoms and protein residues, with colours representing attention weights ranging from 0 (bright) to 1 (dark) (Figure 5(a)). The line plots above and on the right sides of the interaction map represent the mapping of attention scores along the direction of each protein residue or ligand atom, illustrating the distribution of attention across the nodes.

**Figure 5:**
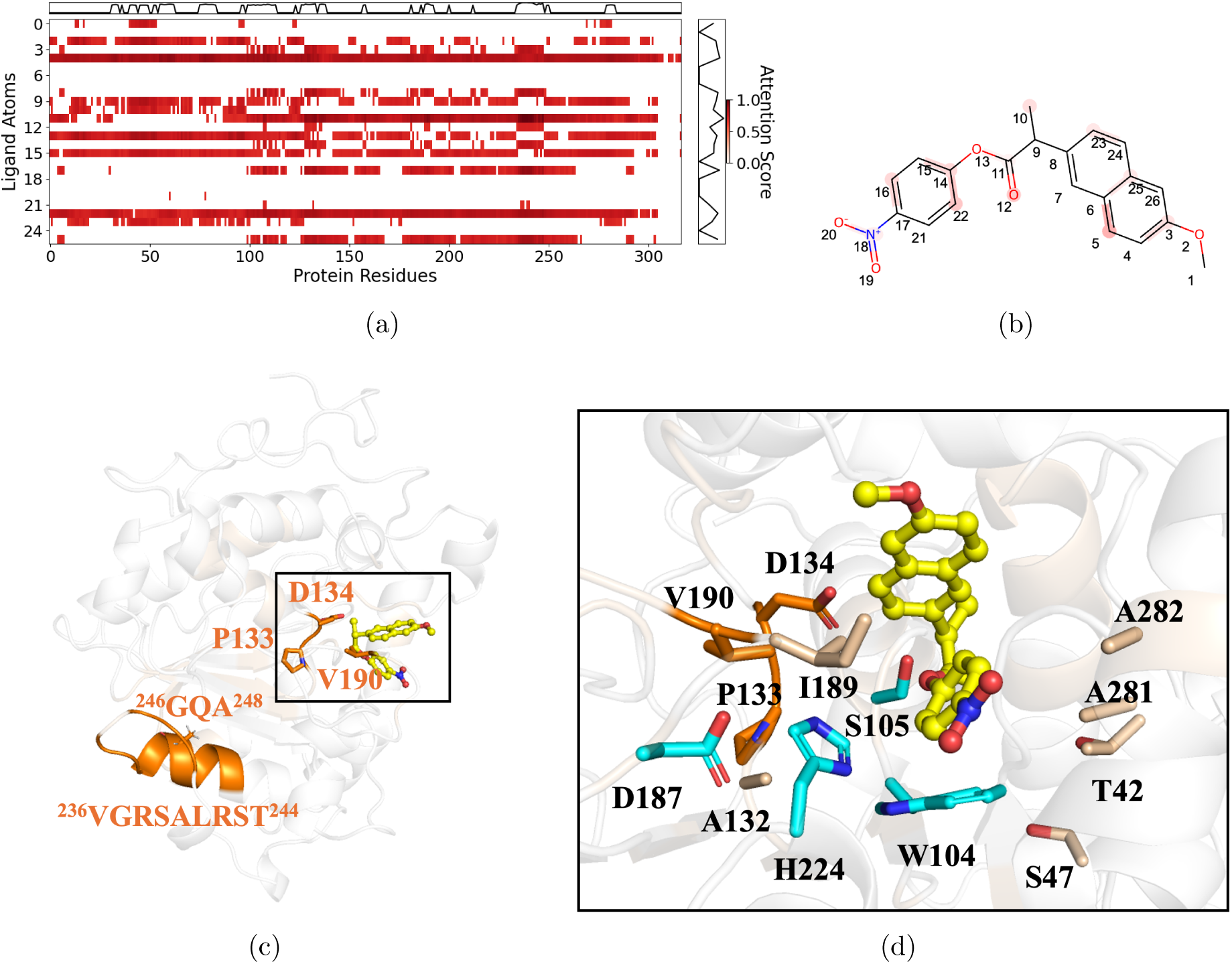
Explainability analysis of the wild-type CalB structure with ligand rac-10. a. Attention heatmaps for protein residues and ligand atoms, where darker colour indicates higher attention weights. Only the points with top 30% of attention weights are retained with their original values, with the others set to 0. Line plots above and on the right of the heatmap show the average attention scores for residues and ligand atoms. b. Structure of ligand rac-10 with attention weights. The ligand atoms with high attention scores (responsible for CPIs) are highlighted in red c. The three-dimensional interaction map of the wild-type CalB protein structure (PDB ID: 1TCA) and the ligand rac-10. The substrate is represented by a ball-and-stick model and marked in yellow. Residues with top 5% attention weights are highlighted in orange colour. d. The binding pose of substrate rac-10. The catalytic triad Asp 187, Ser 105, His 224 and the key residue W104 are marked in cyan colour. The active site residues that are ranked within the top 30% of attention weights are highlighted in pink, while those in the top 5% are marked in orange.

We then compared with the attention weights map with the simulated structure of CalB in complex with rac-10 (Figure 5). The simulated complex was obtained by docking the ligand into the wild-type CalB protein (PDB ID: 1TCA) followed by MD simulations (Supplementary Figure S2). For ligand rac-10, the carbon atoms on its benzene ring show notably high attention weights (Figure 5(b)), indicating that the interactions with the protein residues at these specific sites contributes largely to the overall CPI. This is consistent with the results observed in the MD simulation results, where W104 was found to form a *π*-*π* interaction with the phenyl ring of rac-10 (Figure 5(d)). Previous mutagenesis study showed that mutations at W104 would affect the enzyme’s activity and enantioselectivity.^39^ Interestingly, two second-sphere residue sites T42 and S47 surrounding W104, are also associated with high attention weights. The attention weight importance observed for the third-sphere residues A281 and A282 residues may be attributed to their proximity to the second sphere residue T42. Notably, mutagenesis experiments have shown that the combined mutation at A281 and A282 enhanced Kcat and conversion for ligand rac-10. ^38^ The role of I189 and V190 on CPIs is possibly contributed to their special proximity to another catalytic triad H224. In addition, residues 132-134 (Figure 5(c) and 5(d)) were also found with high attention weights (ranked top 10%). This is possibly because of the hydrophobic interactions between A132 and the key catalytic residue H224. These distantly located residues could affect the enzyme’s catalytic efficiency by indirectly reshaping the active site via the interactions with the first sphere residues. The good agreement between the attention modules in our BioStructNet models and the interaction patterns between protein residues and ligand atoms from the visual inspection, indicate it may serve as a valuable tool to prioritise the important sites for mutation, to improve the efficiency of enzyme engineering.

It should be noted that the residues with high attention weights affecting Kcat and conversion are not limited to the active site. In addition to those residues located in the proximity of the catalytic site, some sites with high attention weights in CPIs predictions are far away from the catalytic site. For instance, we found the residues with the top 5% attention weights are concentrated in the V236-A248 region. Mutation the residues in this region would stabilize the thermostability of this flexible loop, and along with the mutation of other high-weight residues around the catalytic site, would enhance the overall enzymatic activity (Figure 5(c)). Understanding the influence of distantly located residues on enzyme functions is vital for advancements in enzyme engineering. By targeting the modifications of those regions outside the catalytic site, enzyme activity, stability, and specificity may be modulated without directly altering the core catalytic site, resulting in enzyme inactivation. However, experimentally or simulation investigating these effects poses significant challenges which often requires extensive mutagenesis studies and cost. Thus, our BioStructNet model would help to identify the distal residues as candidates for further evaluation, thus significantly reduce the cost of screening large mutant database.

We also examined two CalB variants, RG401 and SG303, which were reported to show distinct enantioselectivity.^38^ Compared with wild-type protein, RG401 has the mutations W104C, L144Y, V149I, V154I, A281C, and A282F; while SG303 has the mutations V149D, I189V, V190C, A281G, and A282V (Figure 6(a)). The attention weights of the RG401 and SG303 variants were compared with that of the WT and the heatmaps were plotted, which reflects the changes of attention weights at each residue position introduced by combined mutations (Figure 6(b)). Red indicates a positive difference in attention weights in the variants compared to the WT, while blue represents a negative difference, with the darker color highlighting regions with notable changes in attention. The heatmap of the RG401 variant displays a significant negative difference in overall attention weights, while the heatmap of the SG303 variant exhibits a more complex pattern with both positive and negative difference at different residue positions. We then compared the structures of these variants in complex with the rac-10 ligand (Figure 6(c)). Residues 189 and 190 exhibit positive attention weight change in both SG303 and RG401 variants, indicating these two positions are responsible for the enhanced yield in the variants compared to the WT. For the RG401 variant, the A281C and A282F mutations caused negative shifts compared with the WT. While in the SG303 variant, the A281G and A282V mutations exhibit positive attention weight shifts. The opposite enantioselectivity of the RG401 and SG303 variants may be attributed to the opposite direction of attention weight shifts of the first and second sphere residues, as demonstrated from the substrate binding environment revealed from high attention weight sites 144, 154, 149, 281, 282 located in proximity to the substrate showed opposite direction of attention weight shifts in the two variants, and mutation of these sites would reshape their substrate binding pockets, such that the substrate exhibits opposite binding orientation in the two variants yielding opposite enantiomer products.

**Figure 6:**
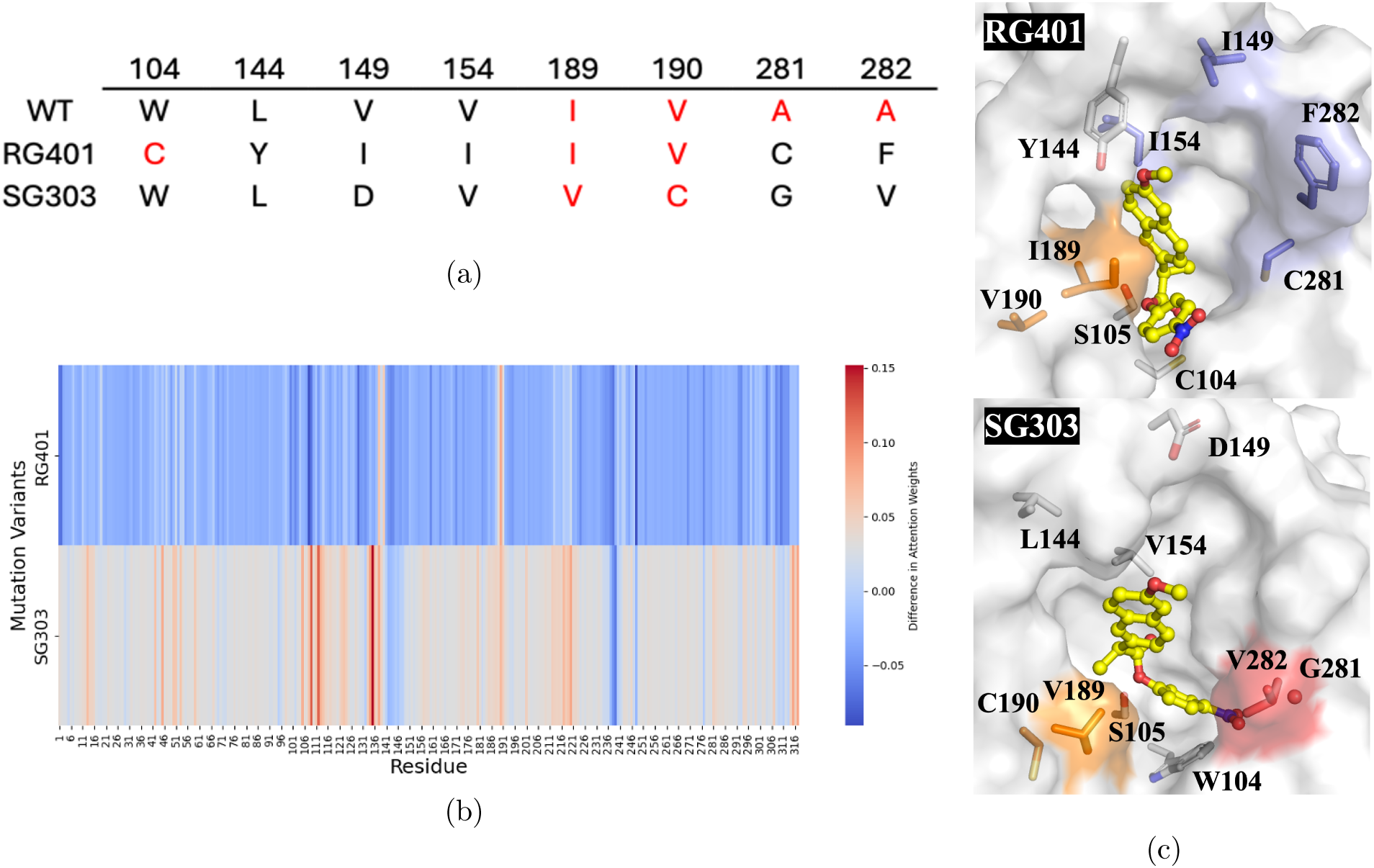
Comparison between two CalB variants RG401 and SG303 with the WT CalB enzyme. a. Mutation sites in the variants. The residues with high attention weight changes compared to the WT are shown in red. b. A comparison heatmap of variants RG401 and SG303 that reflects the differences in attention weights at each site (or residue) between the variants and WT caused by mutations. The colour in the heatmap indicates the magnitude of the difference in attention weights. Colours closer to red (positive difference) or blue (negative difference) signify greater differences in attention weights. c. The three-dimensional structures of the CalB variants RG401 and SG303 in complex with the substrate rac-10. The substrate is represented by a yellow ball-and-stick model. The mutation residues, along with the catalytic residue 105, are coloured based on their respective attention weights predicted by machine learning. Residues with greater attention weight change compared with the WT are shown in red (indicating a positive difference) or blue (indicating a negative difference). Residues V189 and C190, which show a positive difference in both RG401 and SG303, are highlighted in orange.

The comparison of the attention weight of BioStructNet with the simulated complex structure showed that high attention weights align with known critical residues like S105, A281, and A282, while the model also identifies potential distal residues that may influence the enzyme function. The comparative analysis between wild-type and mutated enzymes (RG401 and SG303) further revealed significant differences in attention weights across various residues and the impact of mutations resulting in different protein-ligand binding, thereby validating the model performance in predicting the protein ligand interactions responsible for their functional changes.

## Conclusion

In this study, we developed BioStructNet, a novel deep learning framework tailored for predicting compound-protein interactions (CPIs), with a particular focus on small biocatalyst datasets with specific functions, such as CalB. By leveraging protein’s structural data and advanced graph-based representations, BioStructNet effectively captures both local and global interaction patterns between enzymes and substrates, offering a significant improvement over traditional methods in predicting enzymatic functions.

BioStructNet uses two different interaction modules—Bilinear Co-attention Network (BCN) and a Transformer-based module—to enhance the model’s interpretability and predictive power. The BCN module, with its bilinear attention mechanism, demonstrated its strength in detecting subtle local interactions, particularly useful in processing datasets with high structural similarity. The Transformer-based module, by contrast, provided an insight into long-range and global interactions within protein-ligand complexes through its self-attention and cross-attention mechanisms. The inclusion of layer normalization and a feed-forward network further improved training stability and model performance.

BioStructNet, when combined with transfer learning, significantly outperforms traditional models in predicting enzyme catalytic activities with small datasets. By pre-training the model on large datasets of enzyme-substrate interactions and fine-tuning on small, function-specific datasets, overfitting—a common challenge in small datasets was tackled, while the model’s robustness and accuracy was improved. BioStructNet also enabled to identify critical residues, including those distant from the active site, which may play important roles in enzyme function through allosteric effects or contributions to stabilizing enzyme structures. The CPI prediction in the BioStructNet was validated by molecular docking combined with molecular dynamics simulations. The BioStrucNet provides a comprehensive framework for predicting and understanding protein-ligand interactions. particularly, attention mechanism facilitates the validation of the CPI predictions by simulated ligand protein complex, making it a powerful tool for guiding enzyme engineering and biocatalyst discovery. BioStructNet provides a robust and interpretable method for predicting enzyme-substrate interactions, effectively integrating structural representations and utilizing transfer learning techniques. It would serve as a valuable tool for guiding enzyme engineering efforts, with potential applications in environmental biotechnology and beyond. The successful application of BioStructNet for predicting the functions of small target dataset would lay the ground framework for further model refining and their exploitation across diverse biochemical applications.

## Methods

### Data Collection and Preprocessing

We collected comprehensive databases of enzyme-substrate interactions, encompassing a wide range of enzyme functions and substrates. The data included protein structures, compound structures, and associated interaction properties.

#### Collection and Preparation of Kcat dataset

The source task Kcat dataset for constructing the deep learning model was selectively extracted from the BRENDA^40^ and SABIO-RK^41^ databases using customized scripts through their respective application programming interfaces (APIs). Only enzymes with EC numbers beginning with ’3’ were included, focusing on those involved in hydrolase activities. As the majority of Kcat values reported in BRENDA and SABIO-RK do not specify their assay conditions, such as pH and temperature, these features were not included to maintain the training dataset size and variety. In addition, substrate SMILES, a string notation representing the substrate structure, was extracted by querying the PubChem compound database^42,43^ with substrate names, followed by a python-based script to ensure the canonical SMILES representation of substrates. Several rounds of data cleaning were performed to ensure quality. Protein sequences were queried with two methods: for entries with Uniprot ID information,^44^ the amino acid sequences could be obtained via the application programming interface of the Uniprot with the help of Biopython v.1.78 (https://biopython.org/); and for entries without Uniprot ID, amino acid sequences were acquired from the Uniprot and the BRENDA databases based on their EC number and organism information. The Uniprot ID was used to search the substrate information for each sequence. Subsequently, the UniProt ID was utilized to retrieve the corresponding PDB IDs for each sequence from the ExPASy database,^45^ and only the data with structure information are retained. From the Protein Data Bank,^46^ corresponding PDB IDs are downloaded, and if the structure is a complex, the ligand is deleted, retaining only chain A. Subsequent mutations are modeled using Rosetta^47^ to predict structures, ensuring that if the sequence from the structure file does not match the sequence from UniProt, the sequence from the structure file is used. The final source task dataset, formed for a regression task, consists of Kcat data (in log2 scale) for enzymes with EC numbers beginning with “3” comprising 1664 items with 244 unique PDB structures. The data processing procedure was modified from previous methods,^19,48^ and a detailed explanation of the process diagram, including specific data cleaning steps, can be found in Supplementary Figure S6.

#### Collection and Preparation of Human dataset

Created by Liu et al.,^49^ this dataset includes highly credible negative samples of compound–protein pairs obtained by using a systematic screening framework cotaining 6,728 items. We used a balanced dataset cleaned by previous studies, where the ratio of positive and negative samples was 1:1.^21,35^ BioStructNet requires protein structure information; therefore, we correlate protein sequences with PDB IDs to filter and select data that includes protein structural information. Finally, the human dataset contains 5,568 interactions and 1,726 unique proteins. To compare with benchmark methods on the human dataset, we employed the same partitioning method, utilizing an 80%/10%/10% training/validation/testing random split.

#### Collection, Molecular Dynamics Simulation, and Docking Studies of CalB and Its Variants

The conversion data for substrates by Candida antarctica lipase B (CalB) and its mutations were collected from previous experimental reports.^37,38,50–53^ The CalB dataset supports targeted research focused on optimizing and enhancing this specific activity, which falls under EC 3.1.1.3, belonging to the EC class 3. The dataset contains 233 items with wild-type protein and 65 variants from published literature. SMILE strings of substrates are extracted from ZINC15. ^54^ The three datasets for classification models were generated with three different threshold values, conversion *≥* 15, 30, 40%, respectively. Detailed information about the CalB datasets can be found in Supplementary Figure S8. The cutoff values were selected based on the considerations that: the 15% cutoff helps to distinguish very low conversion rates from moderate and high conversion rates, aiding in identifying cases where enzyme performance is poor or nearly non-functional in substrate conversion; the 30% cutoff establishes an intermediate threshold between moderate and higher conversion rates, providing a midpoint to indicate cases with moderate enzyme activity; the 40% cutoff sets a higher standard to distinguish between moderate and very high conversion rates, emphasizing cases where the enzyme performs exceptionally well. To reflect the mutations effect of CPI compared with wild-type protein, another classification rule is applied by comparing the conversion values of mutants with those of the wild-type protein for the same substrate. If a mutant’s conversion value is comparable to (taking into account experimental errors) or higher than that of the wild-type protein for the same substrate, it is classified as 1; otherwise, it is classified as 0. To enhance the robustness of model training on small datasets and reduce the impact of validation set selection on model performance, we mitigate potential negative impacts from redundant information by focusing on known mechanisms. This is achieved by setting a central point and radius to encircle the pocket area with a spherical region, thereby augmenting the dataset with negative data from outside this pocket. Specifically, the catalytic reaction of the CalB enzyme occurs in an active site pocket controlled by Asp 187, Ser 105, and His 224. These three residues form the catalytic triad, which is central to the catalytic process. We recognized that mutations in the first sphere and second sphere regions around these key residues could impact the enzyme’s catalytic function. However, mutations in residues far from these critical regions (i.e., non-1st sphere and non-2nd sphere mutations) typically have minimal impact on catalytic activity, especially when the distance exceeds 20 Å, as electrostatic interactions and other intermolecular forces are unlikely to effectively influence the active site at such distances. Based on this understanding, we devised a rational data augmentation strategy. The specific steps are as follows: First, using existing substrate information from the database and focusing on 151 residues located beyond the 20 Å range, we randomly generated a series of single-point mutations. These mutations were designed to avoid the catalytic core and secondary interaction regions while ensuring minimal impact on catalytic activity. We generated 1208 such mutation data points and appropriately labelled them as negative samples to reflect their potentially low impact on catalytic function.

The predicted structures of CalB variants were generated using the Rosetta Comparative Modeling (RosettaCM),^47^ leveraging the crystal structure of the wild-type protein (PDB ID: 1TCA)^46^ as a template. For each variant, six candidates generated by RosettaCM were evaluated based on their free energy, and the structures with the lowest free energy were selected to build the molecular dynamics (MD) simulation systems. Each enzyme was solvated in a pre-equilibrated cuboid box of TIP3P^55^ water molecules, ensuring that any protein atom was at least 10 Å from the edge of the box. The system was neutralized by adding Na+ counterions by tleap module in AMBER 20.^56^ A harmonic restraint force constant of 100 kcal mol-1 was applied to the solute molecules and ions to minimize the solvent molecules, followed by 1000 steps of steepest descent and 1000 steps of conjugate gradient unrestrained minimization. A cut-off of 10 Å was used for non-bonded Lennard-Jones potential and electrostatic interactions. Hydrogen bonds were constrained using the SHAKE algorithm during all MD simulations. Progressive heating was performed from 0 K to 300 K over 100 ps (5000 steps with a step size of 0.02 ps) using the NVT ensemble, followed by 1 ns equilibration using the NPT ensemble at 300 K. A harmonic restraint of 5 kcal mol-1 was applied to the solute during equilibration. After equilibration, a 100 ns production MD simulation was conducted using the NPT ensemble at 300 K and 1 bar. We applied cluster analysis to determine the representative protein structures of CalB variants from MD trajectories using the CPPTRAJ module^57^ in AMBER 20. Next, to prepare the functional 3D structure for docking, the represented protein structures were utilized to define the docking box, and the substrate structures were prepared from SMILES using RDKit.^58^ The parameters for model substrate were calculated based on the optimized geometry at B3LYP/6-31G(d) using Gaussian 16. The substrates were docked into the simulated proteins using AutoDock^59^ based on the previously published paper. ^60,61^ The pose corresponding to the highest binding affinity score for each system was selected for subsequent MD simulations. A 100 ns MD simulation was conducted on the enzyme-substrate complex using AMBER 20, which included steps for energy minimization and equilibration. Stable fluctuations in the root mean square deviation (RMSD) indicated that a stable complex structure was achieved. Principal Component Analysis (PCA)^30^ was employed to select the most representative cluster from which the stable structure was extracted.

### Model architecture

In this section, we introduce the framework of structure-based neural network (BioStruct- Net) for predicting CPI of CalB variants by transfer-learned pattern knowledge from large related database, utilizing various data representations and methodologies to address the complexities of biochemical data (Figure 1).

#### GCN for protein structure

The protein structures were represented by contact map graphs, incorporating positional embeddings to maintain spatial relationships. In these protein structure graphs, nodes represent individual amino acids, each characterized by a combination of a one-hot encoding and five key physicochemical properties: molecular weight, *pK_a_*, *pK_b_*, *pK_x_*, and pI. These features offer detailed insights into each amino acid’s mass, acid and base dissociation constants, and isoelectric point, all of which are crucial for understanding their roles in protein structures and interactions. Edges between the nodes signify visual connections based on a distance map and a predefined cutoff 8 Å, ^62,63^ illustrating the spatial proximity of the amino acids within the protein. This module is designed to handle protein data by leveraging the spatial information inherent in protein structures through a combination of positional encoding and graph convolutional networks (GCNs). This approach allows the model to capture both local and global contextual information essential for understanding complex biological functions. The model begins by generating positional encodings for protein nodes, which helps in retaining the sequence order in the otherwise order-agnostic GCN architecture. The positional encoding is mathematically represented by:

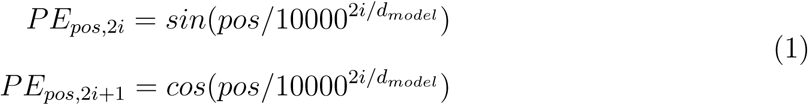

where *pos* is the position and *i* is the dimension. This formulation allows each dimension of the positional encoding to oscillate between different frequencies, ranging from high to low, thereby uniquely encoding each position along the sequence of the protein. Following the addition of positional encodings to the node features, the model applies a series of graph convolutional layers. The GCN layers propagate information across the nodes in the graph. For each layer *l*:

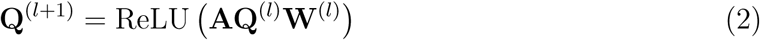

where **A** is the adjacency matrix, **Q**^(^*^l^*^)^ is the node feature matrix at layer *l*, and **W**^(^*^l^*^)^ is the learnable weight matrix. he model sequentially applies three GCN layers, each progressively increasing the feature dimensions to capture more complex patterns in the data. After each convolution, activation functions are applied to introduce non-linear transformations into the model, essential for learning complex functions. The output from the final GCN layer is processed through two fully connected layers to reduce the dimensionality to the desired output size. These layers consolidate features learned from the entire protein graph into a global representation.

#### GCN-GRU for molecular graph

We represented the compound 2D structures as graphs with nodes denoting the individual atoms, and edges denoting the chemical bonds connecting these atoms. Each atom node was represented by its chemical properties, as implemented in the DGL-LifeSci package. Each atom is represented as a 74-dimensional integer vector describing eight pieces of information: the atom type, the atom degree, the number of implicit Hs, formal charge, the number of radical electrons, the atom hybridization, the number of total Hs and whether the atom is aromatic. The combined Graph Convolutional Networks (GCNs) with Gated Recurrent Units (GRUs) module initializes a transformation for the input features to capture both spatial and sequential relationships in ligand. The model begins with an initialization phase where molecular features are transformed into an embedding space. We employed three GNN layers, each was customised combining with a GRU cell to aggregate neighbourhood information for each node:

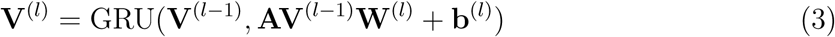

where **A** is the adjacency matrix of the graph, **W**^(^*^l^*^)^ is the weight matrix of the *l−th* layer, **b**^(^*^l^*^)^ is the bias vector of the *l − th* layer, *GRU* is the GRU unit applied to the transformed features and the previous hidden features. The message passing function aggregates messages from neighboring nodes *i* to node *j*:

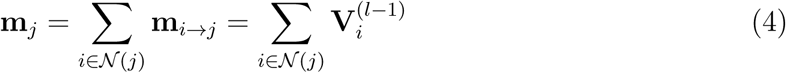

After message passing and aggregation, a linear transformation and RELU activation function are applied. Another GRU update step combines the transformed features with the previous hidden state. Finally, the node features are reshaped to match the batch size and output dimensions.

#### BCN Interaction module

This bilinear attention network was first designed by Bai^16^ to integrate, and process features from two distinct inputs, visual data of ligand and textual data of protein. The BCN module was modified to accept both visual data of ligand and protein with parameters that are configurable to tailor the network’s complexity and regularization. The core of these modules is the bilinear interaction, which can be mathematically represented as:

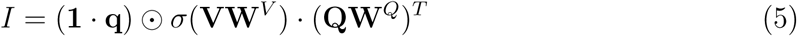

where *V* = *v*^1^*, v*^2^*, …, v^N^* and *Q* = *q*^1^*, q*^2^*, …, q^M^* are the transformed visual features from ligand and protein, respectively, where *N* and *M* denote the number of encoded atoms in a ligand and residues in a protein. *W ^V^* and *W ^Q^* are weight matrices for the *i − th* bilinear interaction. *q* is a learnable weight vector, and 1 is a fixed all-ones vector.I indicates the interaction intensity of respective pairs. denotes the Hadamard (element-wise) product and *σ* denotes the activation function. An elecment I i,j in equation 5 can be written as:

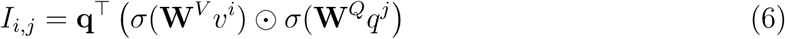

Where: *v^i^* is the *i − th* sub-structural representation of the ligand, and *q^j^* is the *j – th* sub-structural representation of the protein. This bilinear form projects the features into a joint embedding space, facilitating a comprehensive interaction analysis. Following the computation of interaction tensors, the module applies pooling to summarize the spatial information effectively and obtain the *k − th* element of joint representation *f^′^* as:

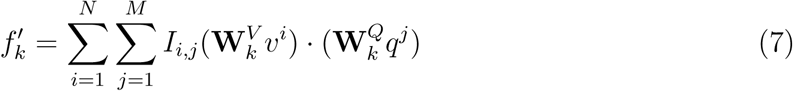

where *W ^V^* and *W ^Q^* denote the *k − th* column of weight matrices. Finally, sum pooling is added on the joint representation vector to obtain a compact feature map:

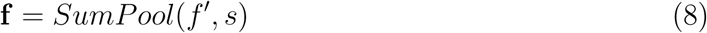

where the SumPool function is a one-dimensional and non-overlapped sum pooling operation with stride s.

#### Transformer-based interaction module

The method encapsulates a neural network architecture that leverages attention mechanisms to process and integrate protein and ligand features. The TransformerBlock first applies self-attention on the protein features to capture dependencies within the protein sequence. This mechanism can be represented as:

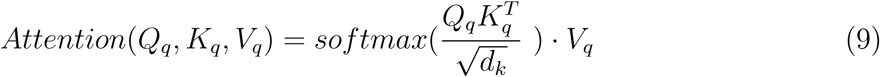

where *Q_q_*, *K_q_*, and *V_q_* represent queries, keys, and values derived from the protein features, respectively, and *d_k_* is the dimensionality of the keys. This operation enhances the representation of protein features by emphasizing informative parts of the sequence. A normalization layer follows this step to stabilize learning by normalizing the layer outputs. Following the self-attention layer, a cross-attention mechanism is employed:

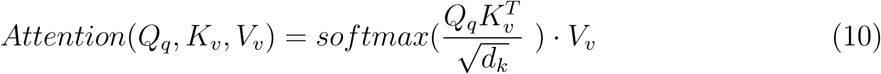

Here, the updated protein features *Q_q_* serve as queries, while both keys *K_v_* and values *V_v_* are derived from the transformed ligand features. This cross-attention step allows the model to dynamically focus on how parts of the protein sequence relate to the ligand features, facilitating a deeper understanding of their interaction. The output of this layer also undergoes normalization.

The outputs from the cross-attention layer are then passed through a feed-forward network (FFN), which typically consists of two linear transformations with a non-linear activation function in between:

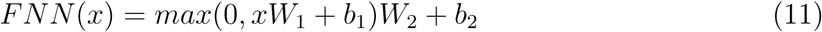

This layer further processes the features to refine the representation for prediction tasks. Finally, a global average pooling operation reduces the feature dimension by averaging across the sequence dimension, which simplifies the output while retaining essential information. The pooled features are then fed into a final linear layer to produce the model’s output, suitable for downstream tasks such as classification or regression.

#### Training and Evaluation

The training process involves pre-training the general model on a large-scale dataset of enzyme-substrate interactions and fine-tuned it on smaller, function-specific datasets to improve prediction accuracy for specific enzyme functions. This transfer learning approach leverages the broad knowledge captured by the general model and adapts it to the specific tasks of interest. For the source task dataset, a 5-fold validation method is employed. This approach is chosen because it allows for a robust estimate of the model’s performance by averaging the results from five different partitions. This helps ensure that the model is tested thoroughly across different subsets of data, providing a reliable measure of its predictive power and generalization ability. For the smaller CalB dataset, bootstrapping is used as the method of validation. Bootstrapping is particularly useful for assessing the stability and generalization ability of models, especially when the sample size is limited. Early stopping is implemented, where training automatically halts if there is no improvement in performance on the validation set for five consecutive epochs. This strategy helps prevent overfitting and conserves computational resources, making it highly effective for managing smaller datasets. To evaluate the performance of the developed models, a comprehensive set of metrics was employed, including Root Mean Square Error (*RMSE*), Coefficient of Determination (*R*^2^), Accuracy, AUC, and Relative Error (RE). *RMSE* is widely used metric for evaluating the accuracy of regression models. It is defined as the square root of the average of the squared differences between the predicted and actual values:

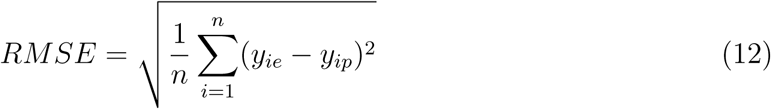

where *y_ip_* is the predicted value, *y_ie_* is the experimental value, *ȳ* is the average of the experimental values and *n* is the total number of items in the dataset (validation dataset or test dataset). *R*^2^ provides a measure of how well future samples are likely to be predicted by the model. It indicates the proportion of the variance in the dependent variable that is predictable from the independent variables.

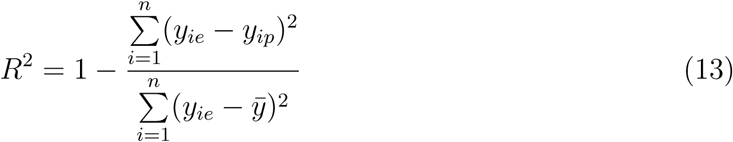

If the predicted responses are sufficiently close to the true values, the *RMSE* would be small. On the contrary, if the predicted and true responses differ substantially the *RMSE* would be large.

Accuracy is the proportion of true results (both true positives and true negatives) among the total number of cases examined, which is a widely used metric to evaluate the performance of binary classification models.:

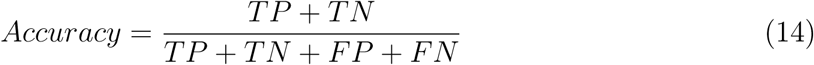

where *TP*, *TN*, *FP*, and *FN* represent the number of true positives, true negatives, false positives, and false negatives, respectively. In the context of classification, RE is used to assess the model’s accuracy in estimating the probability of the positive class. RE for a given threshold is defined as:

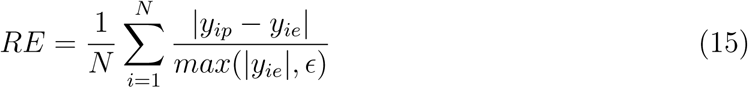

where *y_ip_* is the predicted probability of the positive class, *y_ie_* is the actual class label (0 or 1), *N* is the number of samples, and *ɛ* is a small constant to avoid division by zero. This metric is relevant at 1.0 % probability thresholds to evaluate the high sensitivity of the model’s predictions against changes in the decision boundary. It helps in identifying how well the model’s probability estimates align with true class labels under different operational settings. **Fine-Tuning models.** In this study, we explored the effectiveness of three distinct finetuning methodologies applied to two different interaction modules, BCN and transformer-based interaction modules. This resulted in a total of six unique model configurations, designed to investigate how different fine-tuning strategies influence the performance of interaction modules in capturing complex biochemical interactions. The fine-tuning methods employed are described as follows:

##### Free Layer Fine-Tuning

Unlike the blocked method, free layer fine-tuning allows for the adjustment of all layers within the interaction modules. This method offers the flexibility to adaptively learn from new data across the entire depth of the model architecture, potentially capturing more nuanced patterns and dependencies than more restrictive approaches.

##### Blocked Layer Fine-Tuning

This method involves fine-tuning the output layers within the interaction modules while keeping the remaining layers frozen. By isolating the adjustments to certain layers, this approach aims to refine model responses without extensive retraining of the entire network. It was applied independently to both the BCN and transformer-based modules.

##### LoRa Fine-Tuning

The low-rank (LoRa) fine-tuning method^64^ focuses on optimizing the last layer of the interaction modules, using low-rank matrix adjustments to modify the internal representations within a neural network layer, thereby enhancing model performance while maintaining consistent input and output dimensions. In the transformer-based interaction module, the LoRa fine-tuning method is applied after the attention mechanism. The attention logits are computed solely based on the attention mechanism, and then the adjustments are made to the final layer to refine the model’s output. Similarly, in the BCN interaction module, LoRa fine-tuning is applied after the attention mechanism. However, since the logits in BCN are generated using both the attention outputs and the original lig- and and protein inputs, LoRa adjustments are applied separately to the ligand and protein inputs. This dual adjustment ensures that the fine-tuning process captures the intricate interactions between the ligand and protein more effectively.

### Experimental setting

#### Implementation

BioStructNet is implemented in Python 3.10, along with functions from torch 2.2.2,^65^ Biopython 1.81, ^66^ DGL-LifeSci 0.3.2,^67^ Scikit-learn 1.3.0, ^68^ Numpy 1.26.4^69^and Pandas 2.0.3.^70^ For the initial models, the batch size is set to be 64 and the Adam optimizer is used with a learning rate of 5e-5. We configured the model to use 5-fold cross-validation, with each fold running for a maximum of 100 epochs. For the transfer learning models, we employed a bootstrapping approach with 100 iterations to ensure robust performance estimation. Early stopping was implemented with a patience of 5 epochs to prevent overfitting and conserve computational resources. The best performing model is selected at the epoch giving the best R2 score for regression dataset or accuracy for classification dataset on the validation set, which is then used to evaluate the final performance on the test set. The protein feature encoder consists of three GCN layers with the number of filters [128, 128, 128]. The ligand feature encoder consists of three GCN layers with hidden dimensions [128, 128, 128] as well. In the bilinear attention module, we only employ two attention heads to provide better interpretability. The latent embedding size is set to be 768 and the sum pooling window size is 3. The number of hidden neurons in the fully connected decoder is 512. In the transformer-based interaction module, it is initialized with a model dimension of 128 and utilizes 2 attention heads. The dimension through a feed-forward network (FFN) increases from 128 to 512 and then reduces it back to 128. Our model performance is not sensitive to hyperparameter settings. The configuration details and sensitivity analysis are provided in Supplementary Table S3 and Figure S9.

#### Baselines

We compare BioStructNet with the other models on compound-protein interaction prediction: The shallow machine learning methods — random forest (RF), k-nearest neighbors (KNN) and (L2)—are applied to analyze protein and ligand graph node features aggregated using global sum pooling; Tsubaki’s^35^ and DLKcat^19^ used GNNs to update the atom vectors of molecular graphs, and CNNs to scan the n-gram splited protein sequences, respectively suited for classification and regression tasks; DrugVQA ^21^ is a protein structure-based method employing a dynamic attentive CNN to handle variable-length distance maps of proteins and a self-attentional sequential model to extract semantic features from molecules; TransformerCPI2.0^36^ utilizes TAPE-BERT for protein sequence representation with a self-attention-based transformer encoder and introducies a virtual atom vector to enhance molecular interaction information; ALDELE^20^ is based on five distinct toolkits to integrate sequence-based features for proteins and structure-based physicochemical features of ligands, along with a two-phase attention neural network to predict the interactions; DrugBAN^16^ employs a dual-encoder architecture combining CNNs for protein sequences and GCNs for molecules, which are then combined using a bilinear attention network to capture pairwise local interactions. DrugVQA and DrugBAN were originally designed for classification tasks and have been tailored for the regression tasks here. For the above baseline models, we follow the recommended model hyper-parameter settings described in their original papers.

## Data Availability Statement

Source code, original data and instructions are available at: https://github.com/Xiangwen-Wang/BioStructNet

## Author Contributions

X.W. collected data, developed the model and analyzed the data; X.W. and J.Z. conducted the experiments; X.W., J.Z. and M.H. analyzed the data; M.H. designed and supervised the project. X.W. and M.H. wrote the manuscript. X.W., J.Z, and M.H. interpreted the results and wrote revisions to the manuscript. D.Q. and T.M. contributed to the discussion, editing and approved the final draft.

## Acknowledgement

This work was supported by the Invest Northern Ireland Research and Development Programme, partly financed by the European Regional Development Fund under the Investment for Growth and Jobs programme 2021-2027. The authors are grateful for the computing resources from QUB high performance computing Tier2 computing resource funded by EPSRC (EP/T022175).

