## Supplementary for "BioStructNet: Structure-Based Network with Transfer Learning for Predicting Biocatalyst Functions"

**Table S1** The results of BioStructNet-transformer-model for source dataset.

| Cutoff | TL Models | *AUC* | *Accuracy* | *1.0% RE* |
|---|---|---|---|---|
| 15 | Block | 81.25% | 0.60 | 0.19 |
| | Free | 81.39% | 0.59 | 0.19 |
| | LoRa | 80.79% | 0.67 | 0.24 |
| 30 | Block | 61.86% | 0.56 | 0.42 |
| | Free | 62.66% | 0.57 | 0.37 |
| | LoRa | 62.10% | 0.58 | 0.45 |
| 40 | Block | 59.00% | 0.60 | 0.46 |
| | Free | 54.56% | 0.59 | 0.44 |
| | LoRa | 60.15% | 0.60 | 0.46 |

**Table S2** Experimental results of both conversion and Kcat values for wild-type CalB protein (PDB ID: 1TCA) and variants with different substrates.

| Protein name* | Mutants | Substrate name* | Substrate smiles | Conversion (%) | Kcat (s-1) |
|---|---|---|---|---|---|
| WT | WT | rac-10 | COc1ccc2cc(C(C)C(=O)Oc3ccc([N+](=O)[O-])cc3)ccc2c1 | 9 | 0.05 |
| WT | WT | rac-5 | Cc2ccc(C(C)C(=O)Oc1ccc(N(=O)=O)cc1)cc2 | 39 | 0.18 |
| WT | WT | rac-7 | CC(C(=O)Oc1ccc([N+](=O)[O-])cc1)c1ccc(-c2ccccc2)c(F)c1 | 13 | 0.09 |
| WT | WT | rac-8 | CC(C(=O)Oc1ccc([N+](=O)[O-])cc1)c1cccc(C(=O)c2ccccc2)c1 | 31 | 0.11 |
| WT | WT | rac-12 | CCCCC(CC)C(=O)Oc1ccc([N+](=O)[O-])cc1 | 14 | 0.05 |
| RG401 | WA104C/LA144Y/VA149I/VA154I/AA281C/AA282F/ | rac-10 | COc1ccc2cc(C(C)C(=O)Oc3ccc([N+](=O)[O-])cc3)ccc2c1 | 19 | 0.82 |
| RG401 | WA104C/LA144Y/VA149I/VA154I/AA281C/AA282F/ | rac-12 | CCCCC(CC)C(=O)Oc1ccc([N+](=O)[O-])cc1 | 23 | 4.57 |
| RG401 | WA104C/LA144Y/VA149I/VA154I/AA281C/AA282F/ | rac-5 | Cc2ccc(C(C)C(=O)Oc1ccc(N(=O)=O)cc1)cc2 | 45 | 0.8 |
| RG401 | WA104C/LA144Y/VA149I/VA154I/AA281C/AA282F/ | rac-7 | CC(C(=O)Oc1ccc([N+](=O)[O-])cc1)c1ccc(-c2ccccc2)c(F)c1 | 12 | 0.6 |
| RG401 | WA104C/LA144Y/VA149I/VA154I/AA281C/AA282F/ | rac-8 | CC(C(=O)Oc1ccc([N+](=O)[O-])cc1)c1cccc(C(=O)c2ccccc2)c1 | 6 | 1.66 |
| SG303 | V149D/I189V/V190C/A281G/A282V | rac-10 | COc1ccc2cc(C(C)C(=O)Oc3ccc([N+](=O)[O-])cc3)ccc2c1 | 30 | 0.2 |
| SG303 | V149D/I189V/V190C/A281G/A282V | rac-12 | CCCCC(CC)C(=O)Oc1ccc([N+](=O)[O-])cc1 | 49 | 0.01 |
| SG303 | V149D/I189V/V190C/A281G/A282V | rac-5 | Cc2ccc(C(C)C(=O)Oc1ccc(N(=O)=O)cc1)cc2 | 54 | 0.1 |
| SG303 | V149D/I189V/V190C/A281G/A282V | rac-7 | CC(C(=O)Oc1ccc([N+](=O)[O-])cc1)c1ccc(-c2ccccc2)c(F)c1 | 21 | 0.24 |
| SG303 | V149D/I189V/V190C/A281G/A282V | rac-8 | CC(C(=O)Oc1ccc([N+](=O)[O-])cc1)c1cccc(C(=O)c2ccccc2)c1 | 31 | 0.09 |

* The names of the proteins and substrates are taken from the original papers.

**Table S3** hyperparameter configuration in StructBioNet

| Module | Hyperparameter | Values |
|---|---|---|
| Protein encoder | Protein contact map | Cα |
| | Initial amino acid embedding | 128 |
| | Hidden node dimensions | [128, 128, 128] |
| | Dropout | 0.2 |
| Ligand encoder | Initial atom embedding | 128 |
| | Hidden node dimensions | [128, 128, 128] |
| BCN interaction | Heads of BCN attention | 2 |
| | Sum pooling window size | 3 |
| | Dropout | 0.2 |
| Transformer-based | Heads of self-attention | 2 |
| | Heads of cross-attention | 2 |
| | Hidden dimension | 128 |
| MLP | Hidden dimension | 512 |
| Solver | Learning rate | 5e-5 |
| | Epoch | 100 |

rac-5

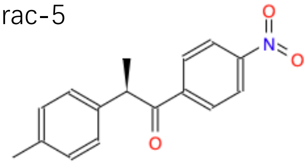

rac-10

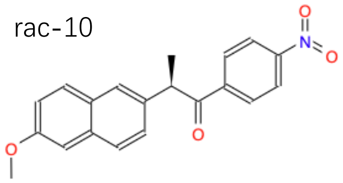

rac-12

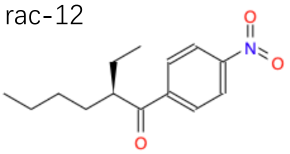

rac-7

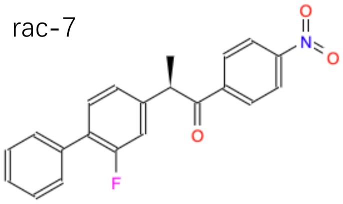

rac-8

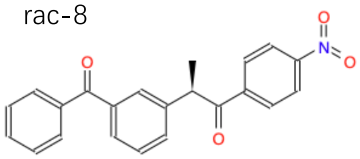

**Figure S1** The molecular structure of “rac-5”, “rac-7”, “rac-8”, “rac-10” and “rac-12”.

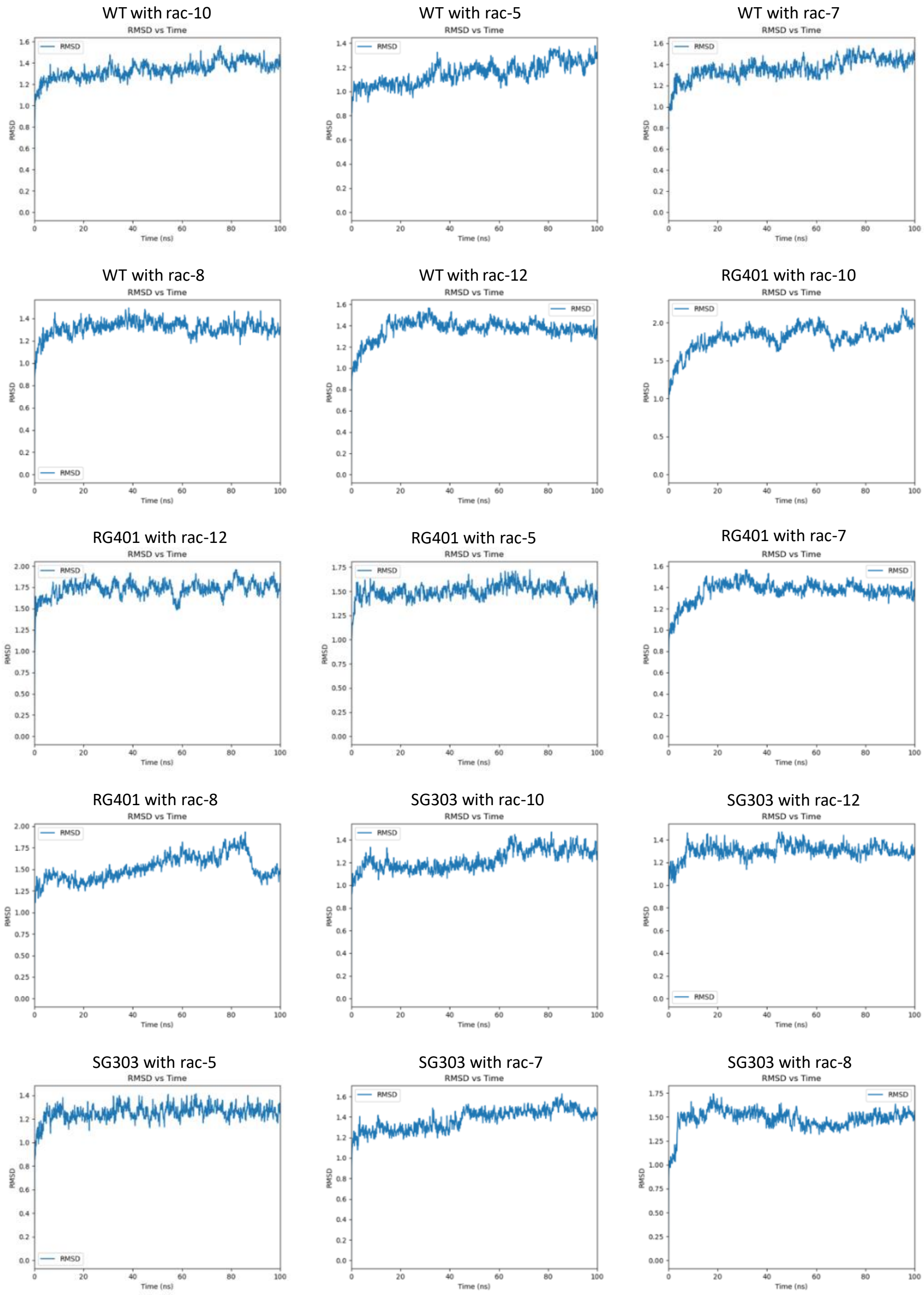

**Figure S2** The RMSD analysis of the MD simulated structures of CalB and its variants in complex with different substrate.

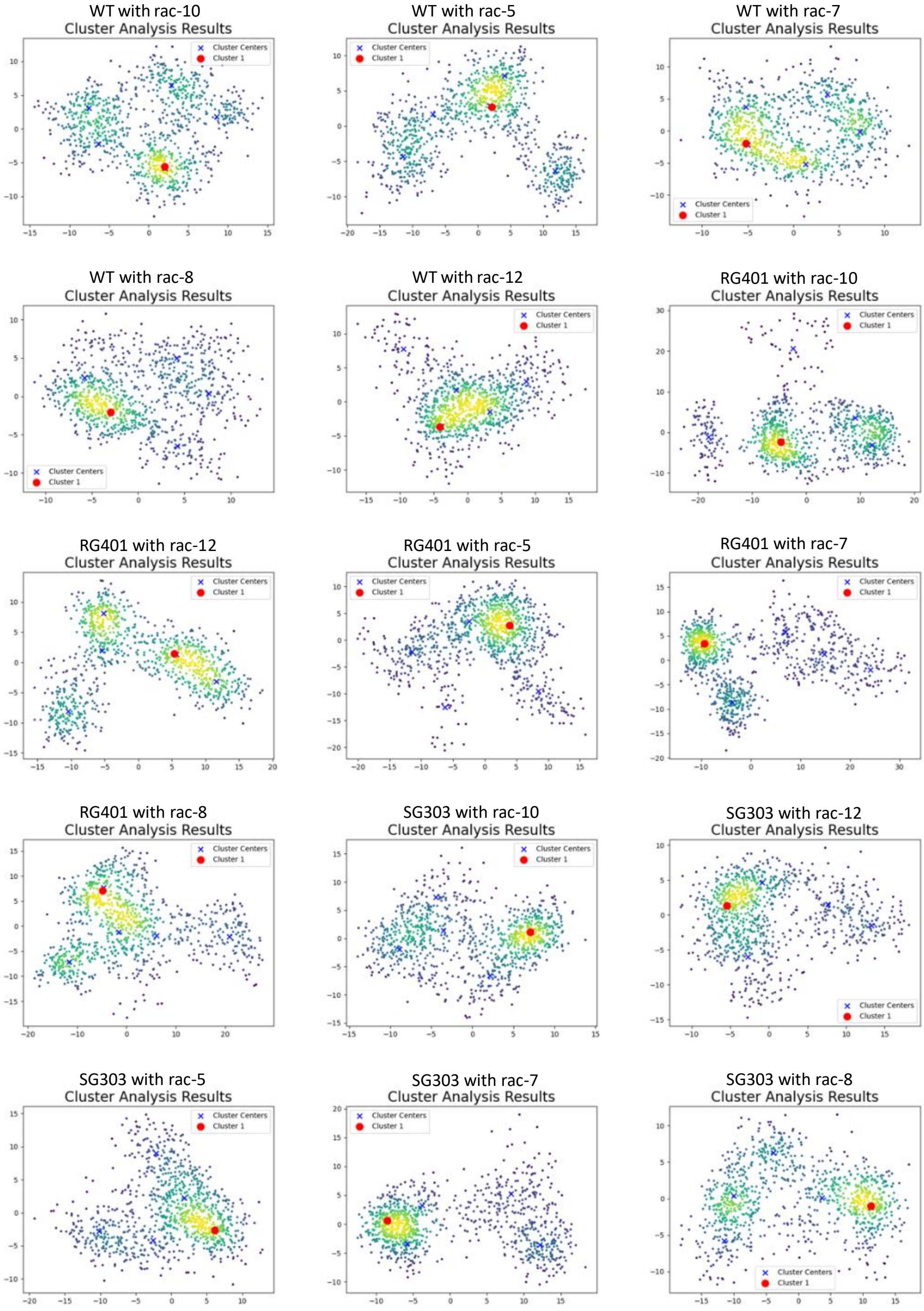

**Figure S3** The conformational distributions of protein−ligand complex ensembles from MD simulations from PCA analysis.

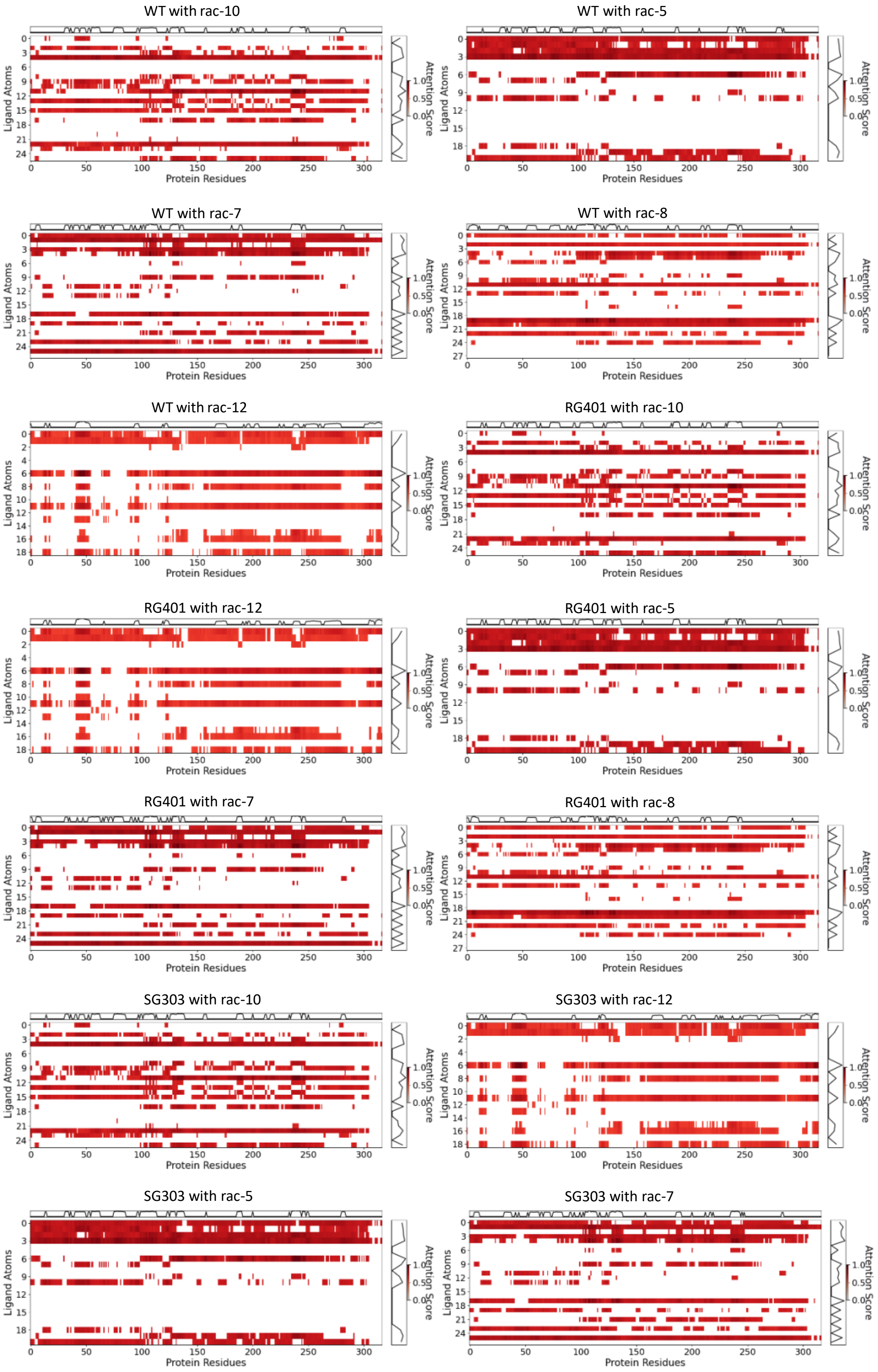
WT with rac-10
WT with rac-5
WT with rac-7
WT with rac-8
WT with rac-12
RG401 with rac-10
RG401 with rac-12
RG401 with rac-5
RG401 with rac-7
RG401 with rac-8
SG303 with rac-10
SG303 with rac-12
SG303 with rac-5
SG303 with rac-7
Ligand Atoms
Protein Residues
Attention Score

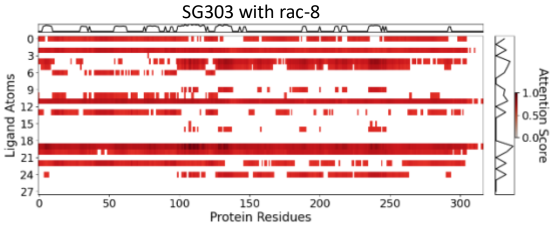

**Figure S4** The machine learning attention heat maps of the CalB-substrate complexes.

WT with rac-10

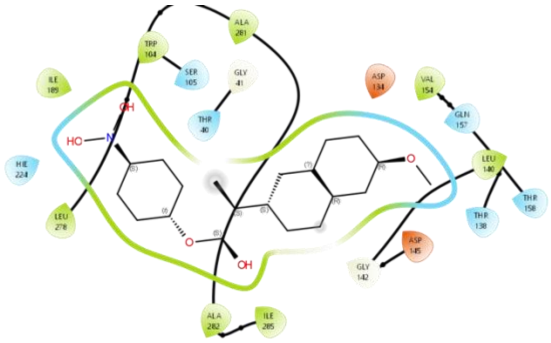

WT with rac-5

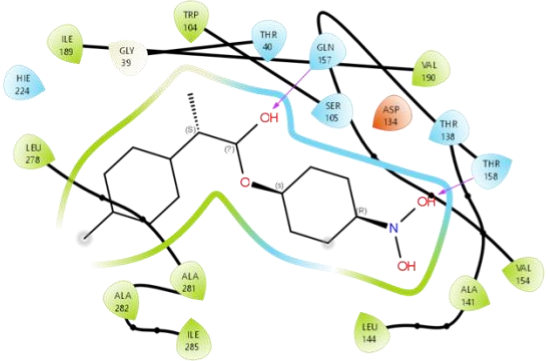

WT with rac-7

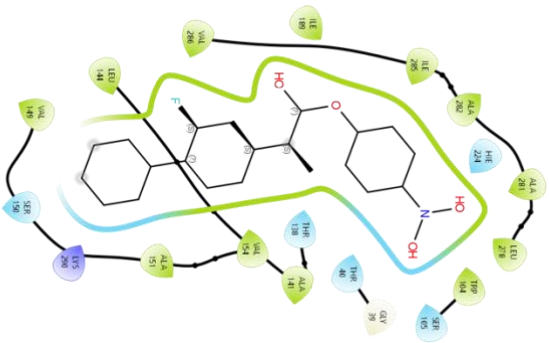

WT with rac-8

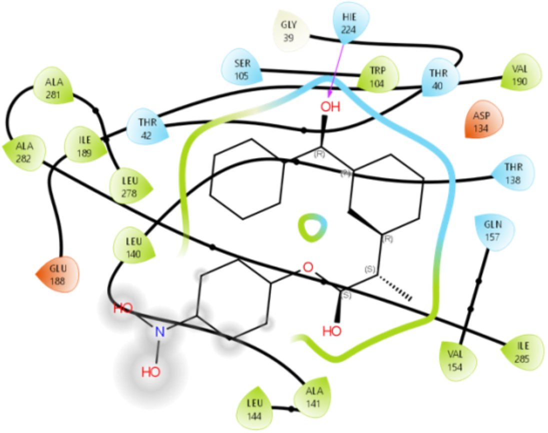

WT with rac-12

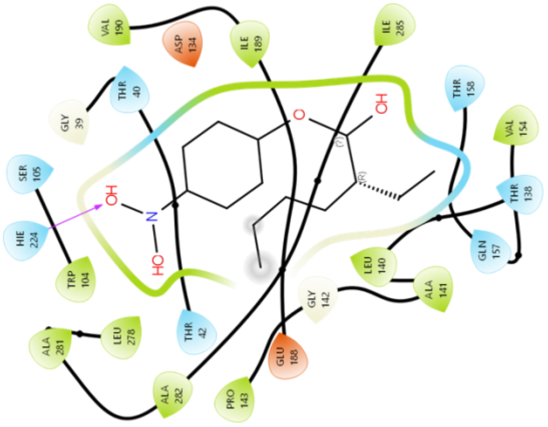

RG401 with rac-10

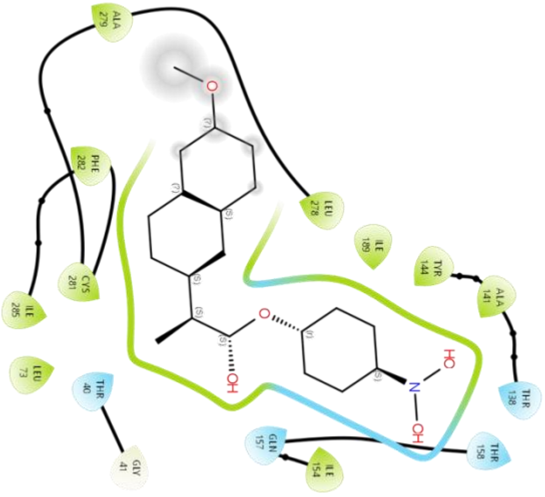

RG401 with rac-12

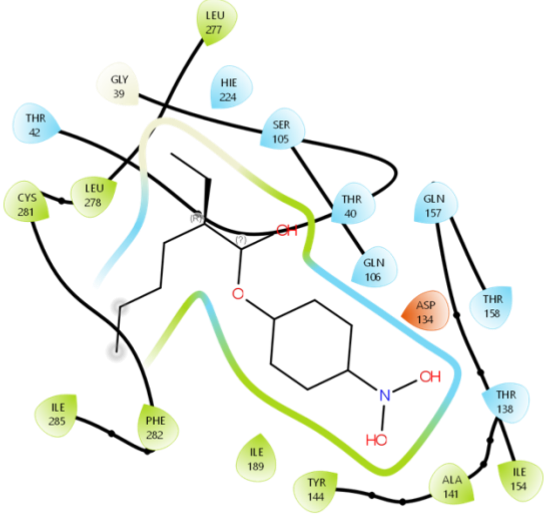

RG401 with rac-5

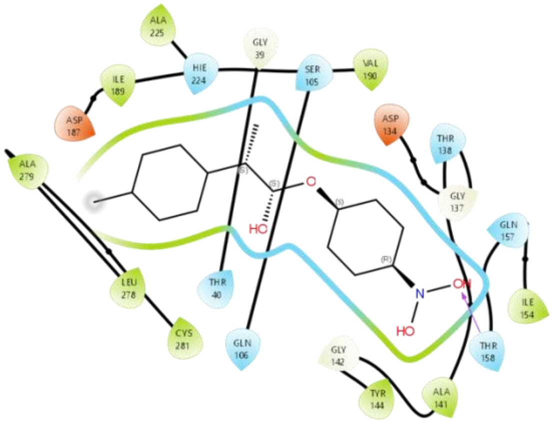

RG401 with rac-7

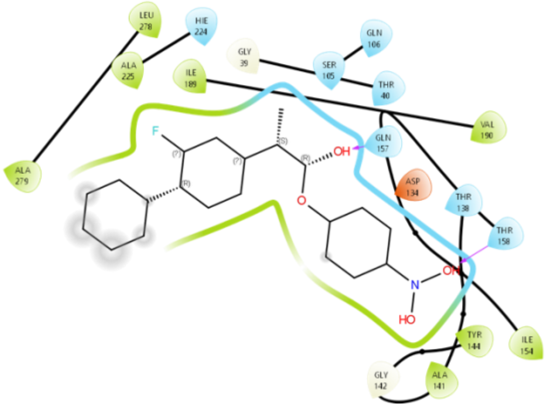

RG401 with rac-8

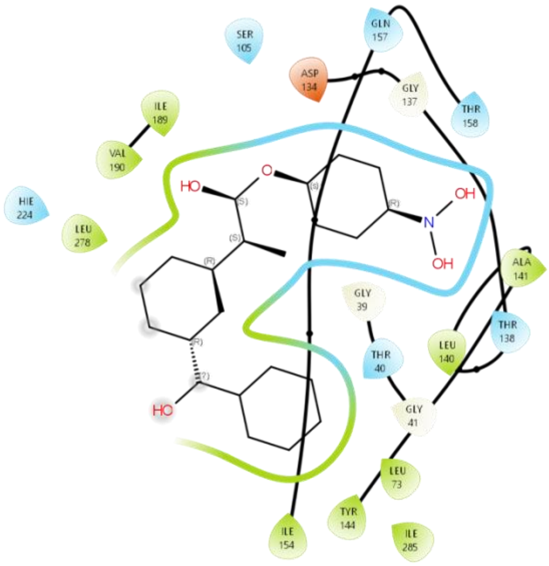

SG303 with rac-10

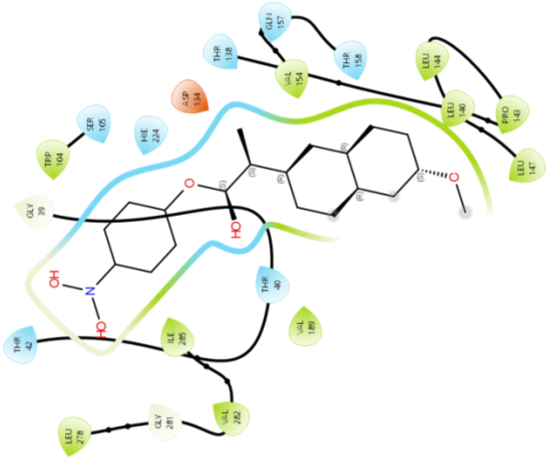

SG303 with rac-12

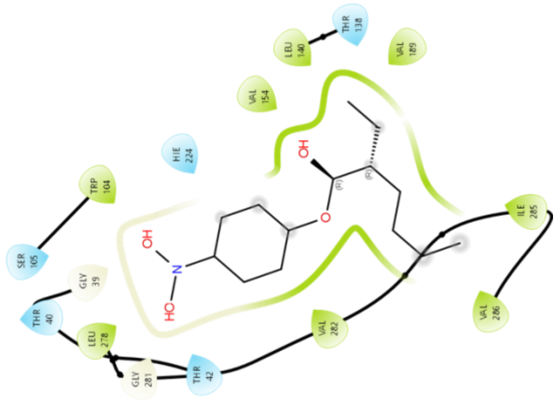

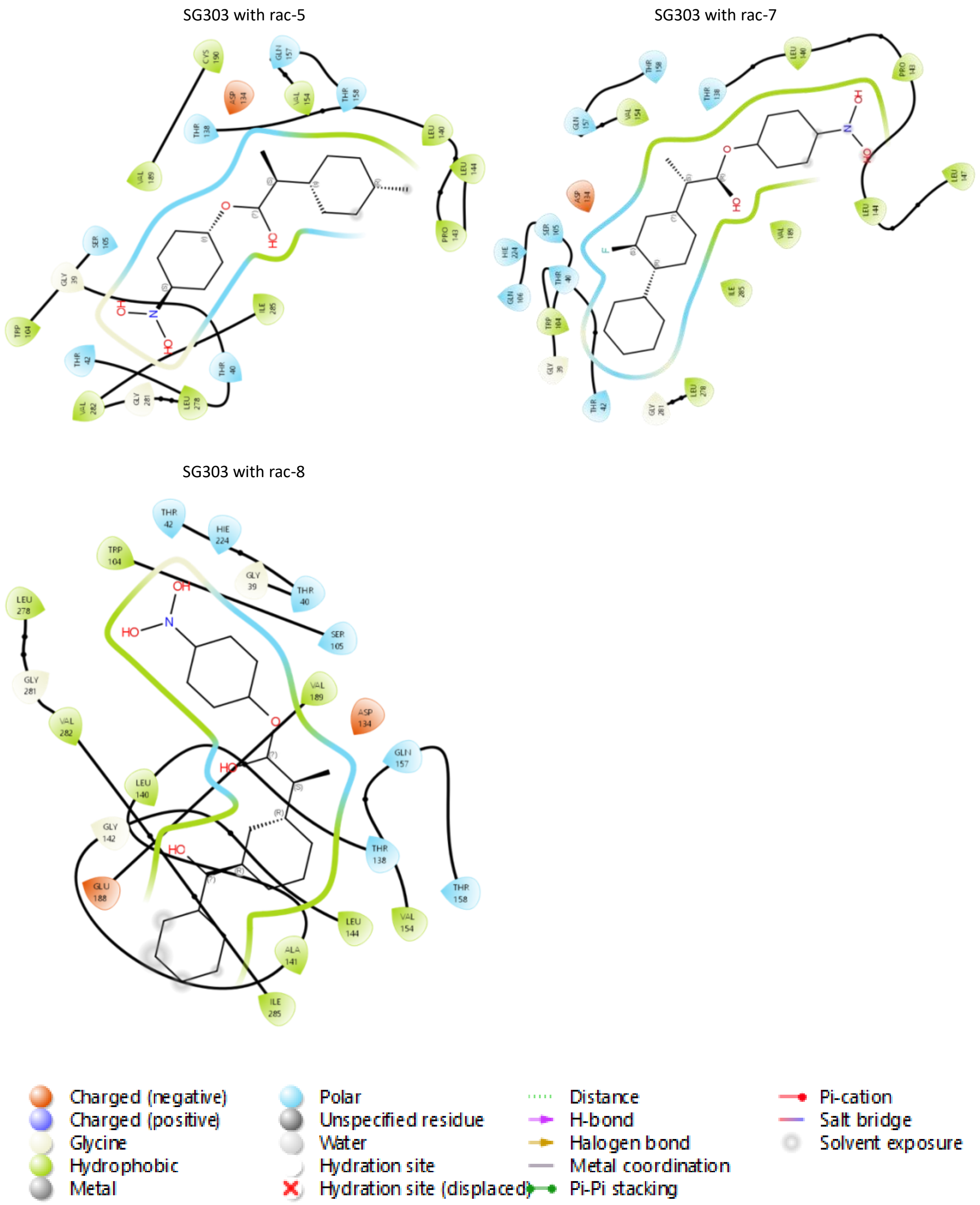

**Figure S5** The 2D interaction maps around the substrate binding pocket in CalB. The complex structures were obtained by molecular docking followed by MD simulations. The graphs are generated by Maestro (Schrodinger Inc).

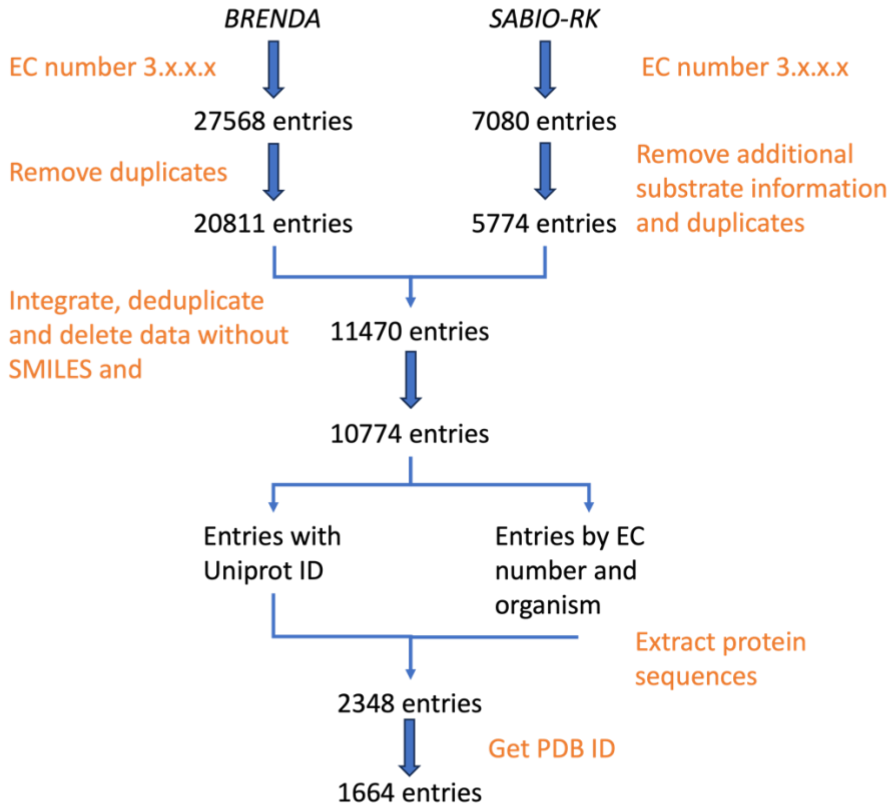

**Figure S6** Data collecting and cleaning for the Kcat dataset for enzymes involved in hydrolase activities (with EC number begin with "3"). The process involves removing duplicates and the irrelevant data, processing additional substrate information and categorizations, extracting protein sequence data, and ultimately obtaining PDB IDs. The scripts and codes are included in Github.

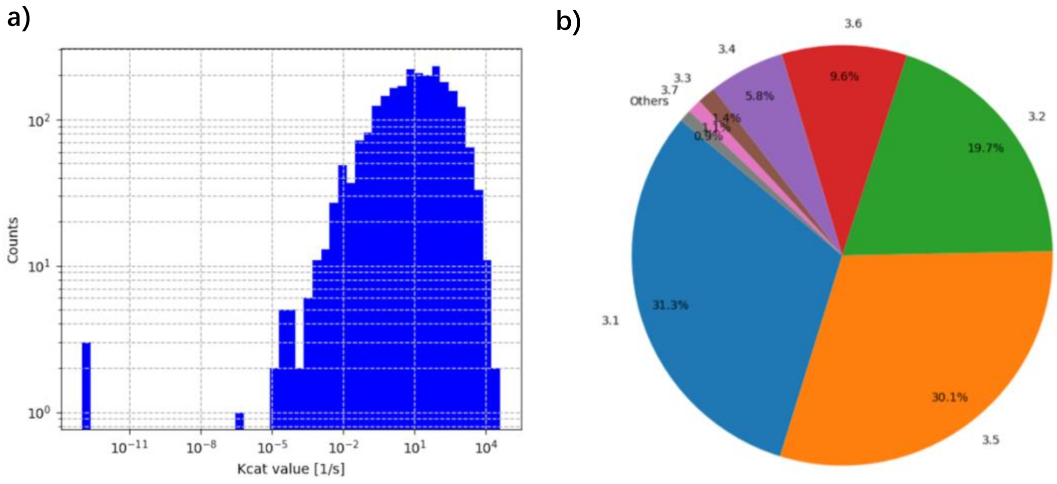

**Figure S7** Distribution of Kcat datasets. a). The distribution of catalytic turnover numbers (kcat) across a dataset on a logarithmic scale. b). The categorization of enzymes based on their EC (Enzyme Commission) number classifications.

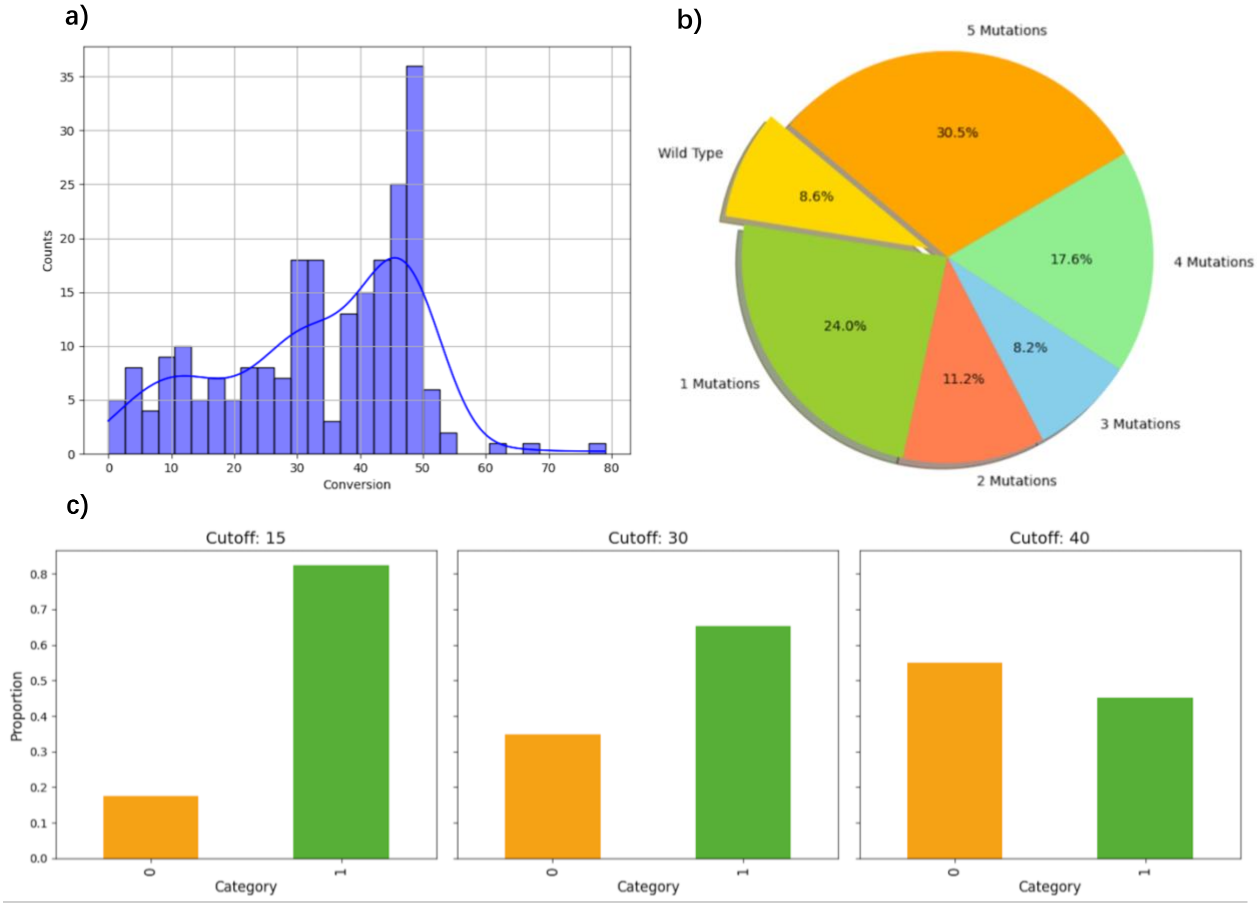

**Figure S8 a).** The distribution of conversion values in the CalB dataset. A kernel density estimate (KDE) is overlaid to indicate the general trend and concentration of data points. **b).** The proportions of wild-type and various mutation counts ranging from one to five mutations. **c).** The distributions of binary categorization of conversion values in the CalB dataset at cutoff thresholds of 15%, 30%, and 40%. “0” indicates lower than cutoff and “1” indicates equal or higher than cutoff.

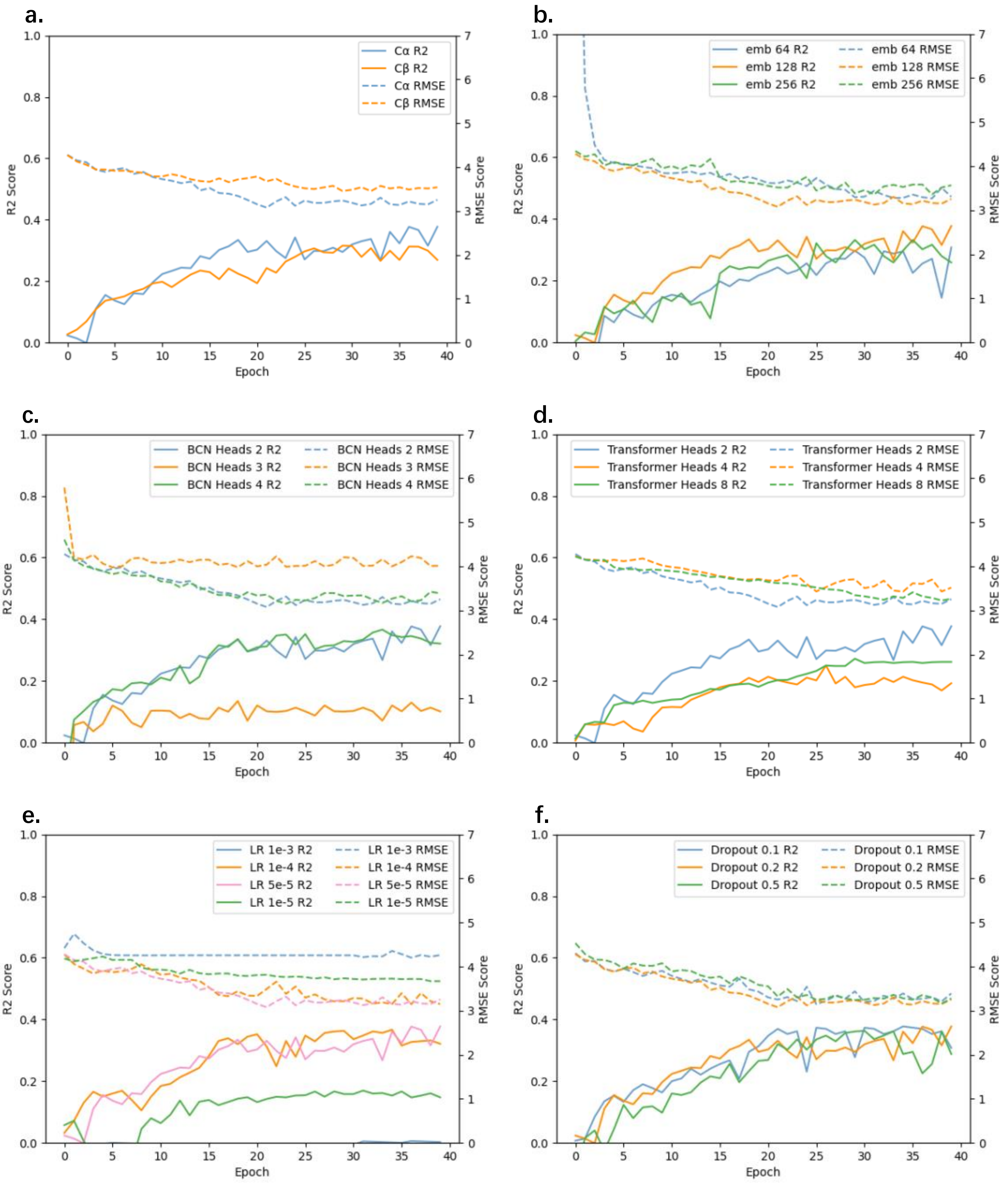

**Figure S9** Learning curves of RMSE and R2 with various hyperparameters on the Kcat validation set. **a).** *Protein contact map*: The choices are Cα and Cβ*, which refer to the types of atoms used to define the contacts between a pair of amino acids in the protein structure. *For most amino acids, the Cβ atom is extracted except for glycine, where the Cα atom is used instead. **b).** *Hidden embedding*: The possible values are 64, 128, and 256. **c).** *attention heads in BCN models*: Available options are 2, 3, and 4. **d).** *attention heads in transformer-based models*: The choices include 2, 4, and 8. **e).** *Learning rate*: Possible values are 1e-3, 1e-4, 5e-5, and 1e-5. **f).** *Dropout*: The options are 0.1, 0.2, and 0.5.
